# Simulations of Pathogenic E1α Variants: Allostery and Impact on Pyruvate Dehydrogenase Complex-E1 Structure and Function

**DOI:** 10.1101/2022.05.17.492373

**Authors:** Hatice Gokcan, Jirair K. Bedoyan, Olexandr Isayev

## Abstract

Pyruvate dehydrogenase complex (PDC) deficiency is a major cause of primary lactic acidemia resulting in high morbidity and mortality, with limited therapeutic options. The E1 component of the mitochondrial multienzyme PDC (PDC-E1) is a symmetric dimer of heterodimers (αβ/α’β’) encoded by the *PDHA1* and *PDHB* genes, with two symmetric active sites each consisting of highly conserved phosphorylation loops A and B. *PDHA1* mutations are responsible for 82-88% of cases. Greater than 85% of E1α residues with disease-causing missense mutations (DMMs) are solvent inaccessible, with ~30% among those involved in subunit-subunit interface contact (SSIC). We performed molecular dynamics simulations of wild-type (WT) PDC-E1 and E1 variants with E1α DMMs at R349 and W185 (residues involved in SSIC), to investigate their impact on human PDC-E1 structure. We evaluated the change in E1 structure and dynamics and examined their implications on E1 function with the specific DMMs. We found that the dynamics of phosphorylation Loop A which is crucial for E1 biological activity, changes with DMMs that are at least about 15 Å away. Because communication is essential for PDC-E1 activity (with alternating active sites), we also investigated the possible communication network within WT PDC-E1 *via* centrality analysis. We observed that DMMs altered/disrupted the communication network of PDC-E1. Collectively, these results indicate allosteric effect in PDC-E1, with implications for development of novel small molecule therapeutics for specific recurrent E1α DMMs such as replacements of R349 responsible for ~10% of PDC deficiency due to E1α DMMs.

## 1. INTRODUCTION

Pyruvate is the end-product of carbohydrate metabolism and after its import into mitochondria by mitochondrial pyruvate carriers is involved in multiple metabolic pathways including gluconeogenesis.^1,2^ The mitochondrial multienzyme pyruvate dehydrogenase complex (PDC) links carbohydrate metabolism to the tricarboxylic acid (TCA) cycle by catalyzing the four-step oxidative decarboxylation of pyruvate to generate acetyl-CoA with concomitant reduction of NAD^+^ to NADH.^3–6^ PDC deficiencies mostly affect the central nervous system and lead to decreased ATP production and energy deficit. ^7–10^

The main components of PDC include the thiamine diphosphate (TPP)-dependent pyruvate dehydrogenase (E1), the dihydrolipoamide acetyltransferase (E2 encoded by *DLAT*), and the NAD-dependent flavoprotein, dihydrolipoamide dehydrogenase (E3 encoded by *DLD*), and E3 binding protein (E3BP encoded by *PDHX*).^3,11^ The inner core of the PDC is formed by up to 60-meric E2 and E3BP structure^12,13^ while lipoyl domain of E2 forms a bridge between E1 and E3^11,14^ that leads to approximately 30 and 12 copies of E1 and E3, respectively^15^. The E1 enzyme is a symmetric dimer of heterodimers (aka heterotetramer; αβ/α’β’) composed of two subunits α (encoded by the X-linked *PDHA1*) and two β (encoded by *PDHB*), that bind two molecules of cofactor TPP in the two active sites, with the catalytic mechanism for activation of TPP and decarboxylation of pyruvate by E1 described.^16,17^ The TPP binding pocket in the E1 heterotetramer consists of α and β subunit residue contacts with some known to be involved in TPP activation and catalytic activity, while the TPP affinity increases in the presence of the substrate pyruvate.^16,18–23^ The irreversible reaction catalyzed by E1 is rate-limiting for the overall activity of PDC^24–26^ and clinically consequential if defective.^10^

Protein-protein interfaces are enriched in tryptophan, tyrosine, and arginine residues.^27^ Buried residues involved in salt bridges (electrostatic interactions at <4 Å), such as arginine, are more likely to be evolutionarily conserved and are more often found at subunit-subunit interface contact (SSIC) forming cooperative networks with other residues stabilizing interface contacts.^28,29^ Greater than 85% of disease-causing missense mutations (DMMs) of E1α are solvent inaccessible (“buried”), and disease-causing replacements of arginine residues constitute 39% of DMMs in E1α.^30^ Furthermore, 30% of solvent inaccessible DMMs of E1α are involved in SSIC.^30^ DMMs of two buried residues in E1α, R349 and W185, destabilize SSICs and result in low PDC activity with clear clinical impact, but the mechanism by which this perturbation is affecting function is not yet clear.^30^

PDC activity is mainly regulated via the phosphorylation of the E1 component with regulation independent of phosphorylation also reported.^11,31–34^ Inactivation of the E1 component of human PDC is achieved *via* phosphorylation of E1α S203, S264, and S271 located on loops formed by E1α amino acid stretches R259-R282 (Loop A, highly conserved) and N195-E205 (Loop B, highly conserved) (Figure 1).^11,31,32,35,36^ E1α S264 is the most rapidly phosphorylated site. However, phosphorylation of any of the three serine residues by pyruvate dehydrogenase kinase (PDK) isoforms is sufficient for inactive E1.^32,35,37^ It was suggested that phosphorylation could regulate PDC activity *via* induction of steric clashes, which result in loop disordering.^17^ Loop A together with Loop B form a wall for the E1 binding site and include residues that are important for the coordination of Mg^+2^ and anchoring of TPP. ^16,17,38,39^ Additionally, active site access and substrate channeling to the lipoyl domain of E2 component can also be controlled by these loops.^17,38,40–42^ It was hypothesized that phosphorylation of E1α S264 would preclude active site access^40^, while computational studies with human E1 components with phosphorylated E1α S264 suggest that the affinity of phosphorylated E1α for pyruvate reduces relative to wild-type (WT) E1α^34^.

**Figure 1.**
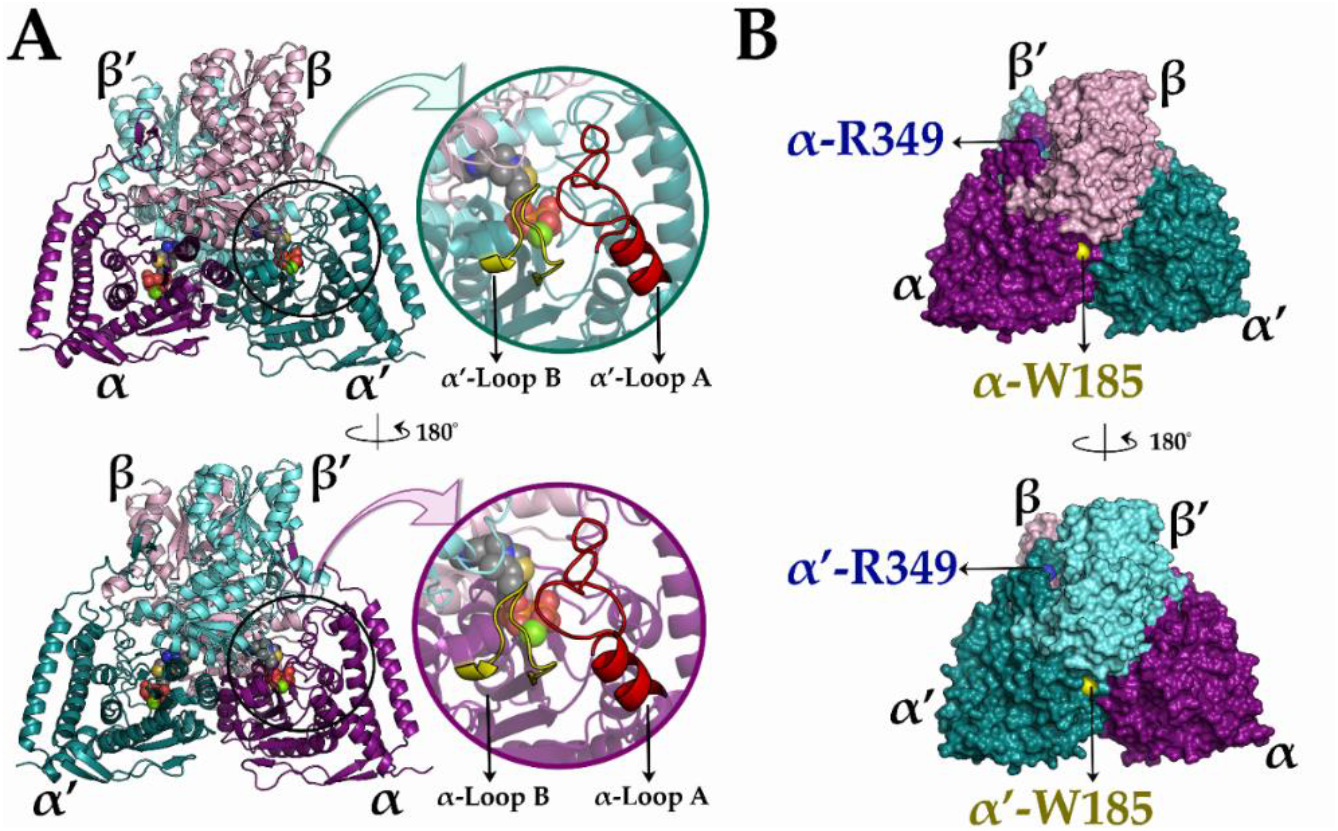
Three-dimensional representations of wild-type (WT) human pyruvate dehydrogenase complex E1 component (PDC-E1). (A) Cartoon representation of heterotetrameric E1 and two phosphorylation loops, Loop A (red) and Loop B (yellow), located on the α (bottom) and α’ (top) subunits. (B) The surface representation of E1 and missense mutation regions, R349 (blue) and W185 (yellow) is located on subunit interfaces.

In this work, we performed molecular dynamics (MD) simulations of WT PDC-E1 and E1 variants with E1α DMMs, R349 and W185, to investigate their impact on human PDC-E1 structure. We evaluated the change in E1 structure and dynamics and examined their possible implications on E1 function with the specific DMMs using backbone mobilities, principal components, clusters, and hydrogen bonds analyses. Because communication is essential for PDC-E1 activity (alternating active sites), we investigated the possible communication network within the WT PDC-E1 *via* centrality analysis using dynamic cross-correlations of residues. The same analyses were performed for E1 variants with E1α DMMs to evaluate their impact on the communication network.

## 2. MATERIALS AND METHODS

### 2.1 System Preparation

Molecular dynamic (MD) simulations of WT E1 and E1 variants with DMMs were carried out starting from the crystal structure of E1 component of human PDC from the protein data bank (PDB ID: 3EXE^17^). This crystal structure contains two heterotetramers, but only one heterotetrameric unit (α_2_β_2_) was selected for MD studies. Males that are affected with PDCD due to DMMs are carriers of replacements of the same amino acid on both the α and α’ subunits, while females affected with PDCD are carriers (heterozygotes) for DMMs. Therefore, simulations for E1 variants with DMMs were performed for samples that were prepared with either a single replacement on only the α subunit (females) or replacements on both the α and α’ subunits (males). This resulted in simulations of nine different samples: WT E1 and E1 variants corresponding to male and female phenotypes.

The protonation states of all amino acids were identified with pKa calculations using pKa-ANI^43^ in conjunction with the reported protonation states in the literature^44^. The systems were described using Ambers ff14SB force field parameters for proteins.^45^ The coordinate and the topology files of the samples were prepared after the hydrogenation of the samples using the *tLEaP* module of AmberTools21.^46^ The force field parameters for TPP were prepared using Antechamber program of AmberTools21^46,47^ while the partial charges were obtained from literature^44^. The systems were modeled with a Mg^+2^ metal ion that anchors the diphosphate group of TPP that has the net charge of −2. The waters in the crystallographic structure were removed and samples were solvated using OPC3 water model^48^ with a 12 Å minimum distance between the solute and simulation cell edges resulting in an average box dimension of 128 Å x 134 Å x 110 Å. Samples were neutralized by the addition of 8 Na^+^ ions for WT E1 and E1 variants with W185C replacements, 9 Na^+^ ions for E1 variants with R349 replacements on only the α subunit, 10 Na^+^ for E1 variant with R349 replacements on both the α and α’ subunits, and 7 and 6 Na^+^ ions for E1 variants with W185R replacement on only the α subunit and W185R replacement on both the α and α’ subunits, respectively. The salt concentration was provided with additional 5 Na^+^ and 5 Cl^-^ ions.

### 2.2 Molecular Dynamic Simulations

An 11-step equilibration procedure^49^ that consists of harmonic restraints on the protein residues and its reduction in each step at 10 K, followed by the gradual heating of samples to 300 K with a gradual harmonic restrain reduction at 300 K, was performed for each sample prior to production simulations using AMBER20’s *pmemd* module.^46^ Each production simulation of 500 ns was performed in the NVT ensemble at 300K using the CUDA-accelerated version of AMBER20’s *pmemd* module.^46,50,51^ A timestep of 2.0 fs was used along with the Berendsen temperature coupling scheme^52^ and the SHAKE algorithm^53^ for the bonds involving hydrogen atoms. Long-range electrostatics were computed with the particle mesh Ewald summation (PME) technique^54^ using a real-space cutoff distance of 8 Å.

### 2.3 Trajectory Analysis and Visualizations

Calculations of root mean square deviations (RMSD), root mean square fluctuations (RMSF) of the residues, dynamic cross-correlation matrices, and any hydrogen bond, principal components and clustering analyses of trajectories were performed using the *cpptraj* module of AmberTools21.^46,55^ The betweenness and eigenvector centralities were computed with dynamic cross-correlation matrices of the trajectories using *correlationplus*^56^ Python package. The clustering analyses were performed using the hierarchical agglomerative (bottom-up) approach. The distance between the frames was computed via the best-fit coordinate RMSD using the C^α^ atoms on the phosphorylation loops, and clustering was finalized when the minimum distance between the clusters was greater than 2.0 Å. All three-dimensional representations were obtained using VMD^57^ and Schrödinger’s Pymol^58^.

## 3. RESULTS AND DISCUSSION

The impact of DMMs on E1 dynamics was evaluated with simulations of WT E1 (Figure 1) and E1 samples with DMMs (referred here as E1 variants with DMMs). Simulation stability along the trajectories was determined with root mean squared deviation (RMSD) of backbone atoms (C, C^α^, and N) with the initial structure of the variant as a reference for the corresponding sample. The RMSD values of all the samples were below 2 Å (Supplementary Figures S1 and S2), indicating sufficient stability for further analysis.

### 3.1 Effect of E1α DMMs on Mobility

Mobility of residues were evaluated by root mean squared fluctuations (RMSF) of C^α^ atoms. To assess the impact of DMMs on mobility, relative RMSF values *(ΔRMSF)* of each E1 variant with the specific E1α DMM were computed relative to WT E1 (Figure 2). We found that the relative mobility of all the evaluated E1 variants stayed within ±1 Å for almost all residues (Figure 2). However, significant mobility losses or gains were observed for residues within the phosphorylation loops (Loop A, representing amino acid stretch R259-R282) on both the α and α’ subunits of E1 (Figure 2).

**Figure 2.**
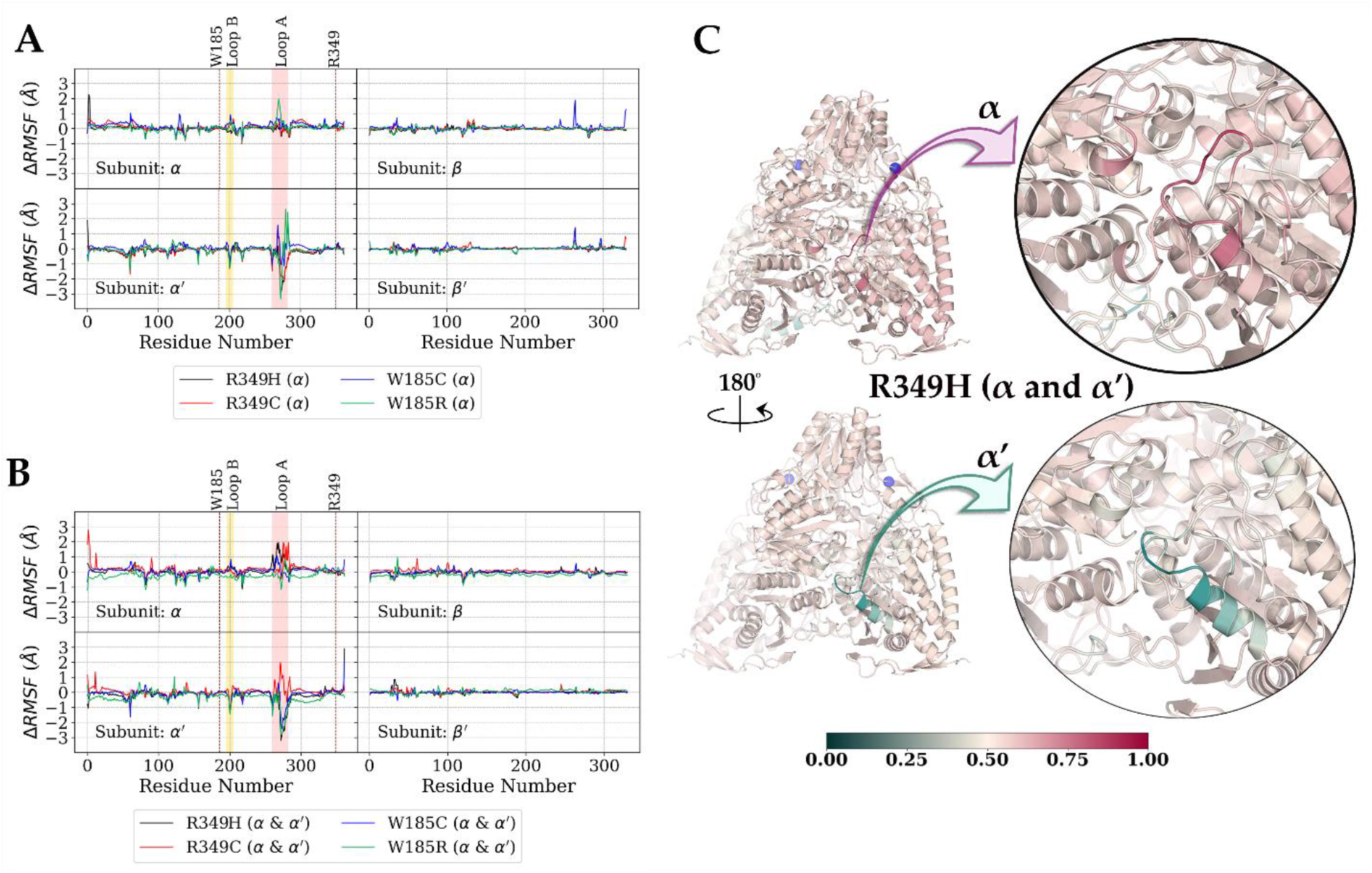
Root mean squared fluctuations (RMSF) values of C^α^ atoms on (A) variants with a mutation on the α subunit relative to wild-type (WT), and (B) E1 variants with mutations on both the α and α’ subunits relative to WT indicate that mutations can drastically affect the dynamics of Loop A on the α and α’ subunits. (C) Three-dimensional representation of the average simulation structure of variant R349H (α and α’) colored by normalized relative RMSF values of C^α^ atoms. Mutation (R349H) locations are depicted with blue sphere.

#### 3.1.1 Amino acid replacements on the a subunit only

A single amino acid replacement on a single α subunit represents a situation observable in females who are carriers (heterozygotes) for a pathogenic missense X-linked *PDHA1* mutation and affected with PDCD. For E1 variants with a DMM only on the α subunit, the major difference in ΔRMSF was at or near Loop A of the α’ subunit (Figure 2A). Replacements of E1α R349 (R349C and R349H) resulted in a loss of mobility of their respective α’-Loop A, particularly in the amino acid stretch S271-E277 (Figure 2A and Supplementary Figures S3A and S4A). The loss of mobility with E1α R349H replacement was higher than that observed with E1α R349C replacement (Figure 2A). With E1α W185 replacements (W185C and W185R), either an increase or a decrease in mobility was observed depending on the region of the α’-Loop A (Figure 2A and Supplementary Figures S5A and S6A). However, with either of the E1α W185 replacements, the α’-Loop A end became much more mobile relative to WT E1 α’-Loop A (Figure 2A). A decrease in mobility was observed with E1α W185R in the amino acid stretch G269-T274 within α’-Loop A (Figure 2A). Interestingly, E1α W185R was observed to be the only variant tested that had mobility changes of up to about +2 Å on α-Loop A (Figure 2A). These results show that the type of amino acid replacement at a specific E1α region could have different E1 dynamic consequences at Loop A, which is consistent with the clinical variability noted in females with different replacements of specific E1α amino acids.^30,59^

#### 3.1.2 Amino acid replacements on both the α and α’ subunits

Replacement of the same amino acid on both the α and α’ subunits represents a situation observed in males with a single chromosome X and affected with PDCD due to a pathogenic missense *PDHA1* mutation. Not surprisingly, E1 variants with DMMs on both the α and α’ subunits showed different mobility changes compared to E1 variants with a single DMM on the α subunit (compare Figure 2A and 2B). For E1 variants with R349 replacements (R349C and R349H) on both the α and α’ subunits, an increase in the mobility relative to WT E1 was observed for different regions of α-Loop A (Figure 2B and 2C and Supplementary Figure S3B). The mobility of the α-Loop A with W185 replacements (W185C and W185R) on both α and α’ subunits was almost similar to WT E1 (mobility changes of ±1 Å; Supplementary Figures S5B and S6B). In contrast, α’-Loop A lost its mobility relative to WT E1 for R349H (Figure 2C), W185C (Figure 2C and Supplementary Figure S5B), and W185R (Figure 2C and Supplementary Figure S6B) replacements on both the α and α’ subunits, particularly in the amino acid stretch S271-E280. With R349C replacement on both α and α’, the mobility of α’-Loop A remained within ±1 Å except for residues S271 and Y272 with Δ*RMSF1* 1.5 Å. These results show that not only does a DMM on one (Section 3.1.1) or both α subunits alter E1 dynamics, but also the same residue replacement on both the α and α’ subunits affects the phosphorylation loops differently depending on the nature of replacement.

Changes in the mobility of phosphorylation Loop A can have multiple consequences. First, Loop A contains residues that anchor the cofactor TPP. E1α H263 interacts with the diphosphate tail of TPP, and its replacement causes a near-zero PDC activity.^17,39,41^ This residue is also in close proximity to E1α H63, Y89, and R90, all of which interact with the phosphate groups of TPP.^16,30,39^ Second, Loop A residues are in close proximity to regions important for the coordination of Mg^+2^. Besides the phosphate groups of TPP anchored by Loop A residues, E1α R259, Y272, and E277 are packed against E1α Y198, which coordinates Mg^+2^ via its backbone.^17,39^ Therefore, any change in mobility of Loop A could have a direct effect on the Mg^+2^ coordination which can hinder the catalytic activity. It should also be noted that in *Escherichia coli* PDC, the dynamics of the homologous loops were reported to control the rate-limiting step (formation of C2α-lactylthiamin diphosphate).^60^ Third, Loop A is involved in substrate intake or acetyl transfer. For instance, conformational changes of E1α H271 in *Geobacillus stearothermophilus* (homologous to E1α H263 in human) are proposed to be involved with the transfer of acetyl group from TPP to the lipoyl domain of E2.^42^ The open and close movements of loops in *E. coli* E1 is important for E2 lipoyl domain binding and E1-E2 active center communication.^61–63^ Furthermore, opening/closing of the β domain loop in TPP-dependent yeast pyruvate decarboxylase is reported to be crucial for product release, which is the rate-limiting step of the catalysis.^64,65^ Therefore, Loop A mobility changes may affect the active site reaction by disrupting the TPP anchors and/or coordination of the Mg^+2^ ion as well as proper substrate intake and/or transfer of the acetyl group to the lipoyl domain of E2.

### 3.2 Principal Components and Clusters Analyses

The dynamics of WT E1 and E1 variants with DMMs on one or both α subunits were investigated with principal component analysis (PCA) (Figure 3 and Supplementary Figures S7-S20), while different conformational states along the trajectories were studied with the representative structures of highly populated clusters (Figure 4 and Supplementary Figures S7-S20). All E1 variants, including WT E1 showed functional motions at the phosphorylation loops A and B. The WT E1 essential motion, particularly on the α’ subunit, is the opening and closing of Loop A (Figure 3A left panel; states R and P, respectively). This motion might enable access of the substrate or the lipoyl group of E2 to the E1 active sites in a similar fashion as was reported for *E. coli* PDC.^61,62^ Clustering analysis of the WT E1 yielded a highly populated conformational state with closed α’-Loop A (50% of the trajectory, state P; Figure 4A left panel), while the same loop was open in the second most densely populated cluster (20% of the trajectory, state R; Figure 4A right panel). These observations, in conjunction with the centrality analysis (see Section 3.3 below) support the proposed shuttle like mechanism which states that a channel at each access site opens and closes with the help of a flip-flop action.^16^

**Figure 3.**
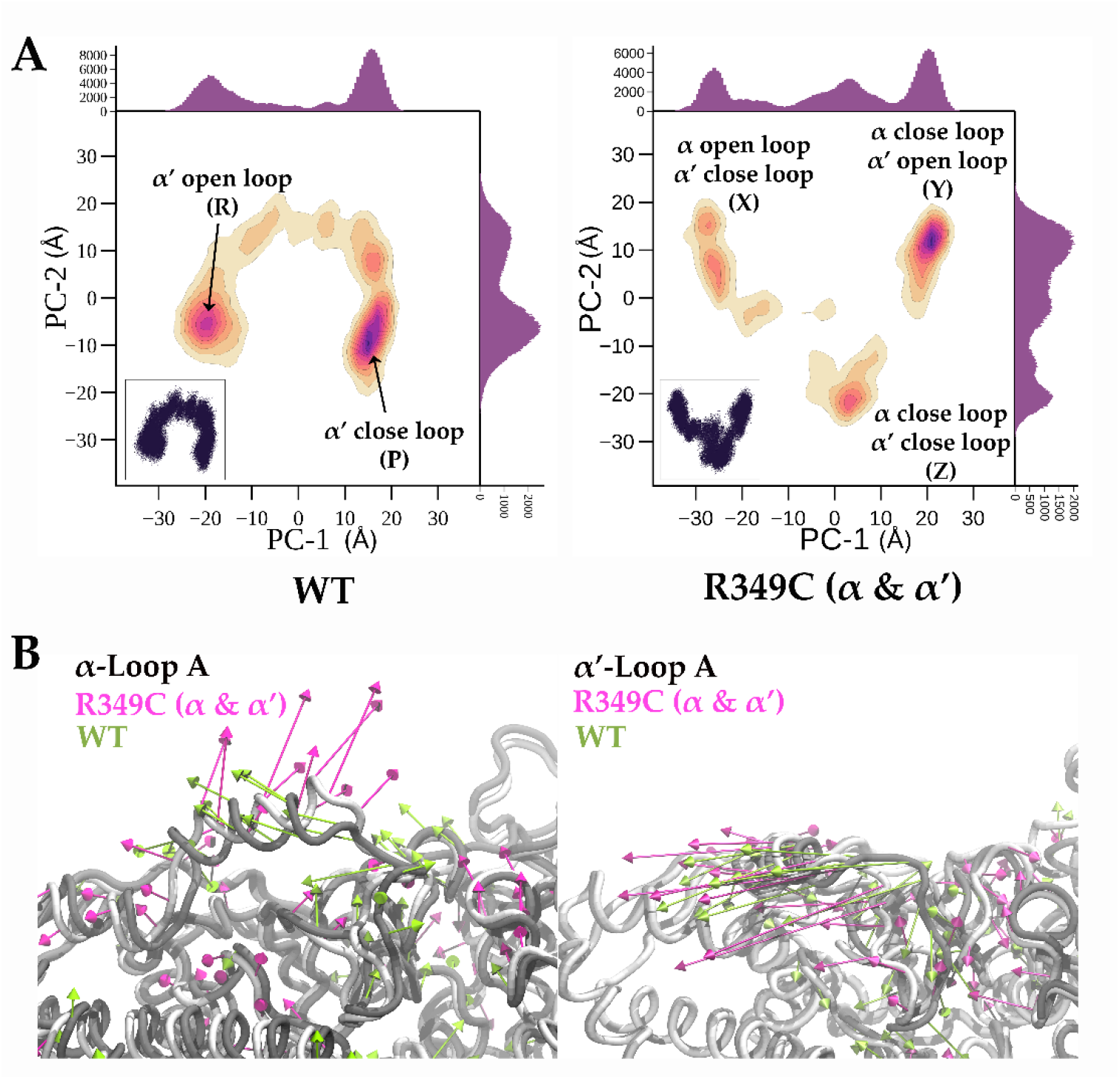
Principal component analysis (PCA) provides more insights on the dynamical properties of wild-type (WT) E1 and E1 variants with missense mutations. (A) Two-dimensional density plots for trajectories of WT E1 (left) and E1 variant with R349C replacement on both the α and α’ subunits projected onto the subspace spanned by the first two principal components, PC-1 and PC-1. Scatter plots for the same components are shown lower left corners of density plots. The populations are denoted with 2D histograms at the top (PC-1) and right (PC-2) of density plots. The regions with a high population on density plots correspond to structures with open/close Loop A. (B) First principal components (PC-1) of Loop A on α (left) and α’ (right) subunits showing alterations of E1 variant with R349C replacement on both the α and α’ subunits (magenta) compared to WT E1 (green).

**Figure 4.**
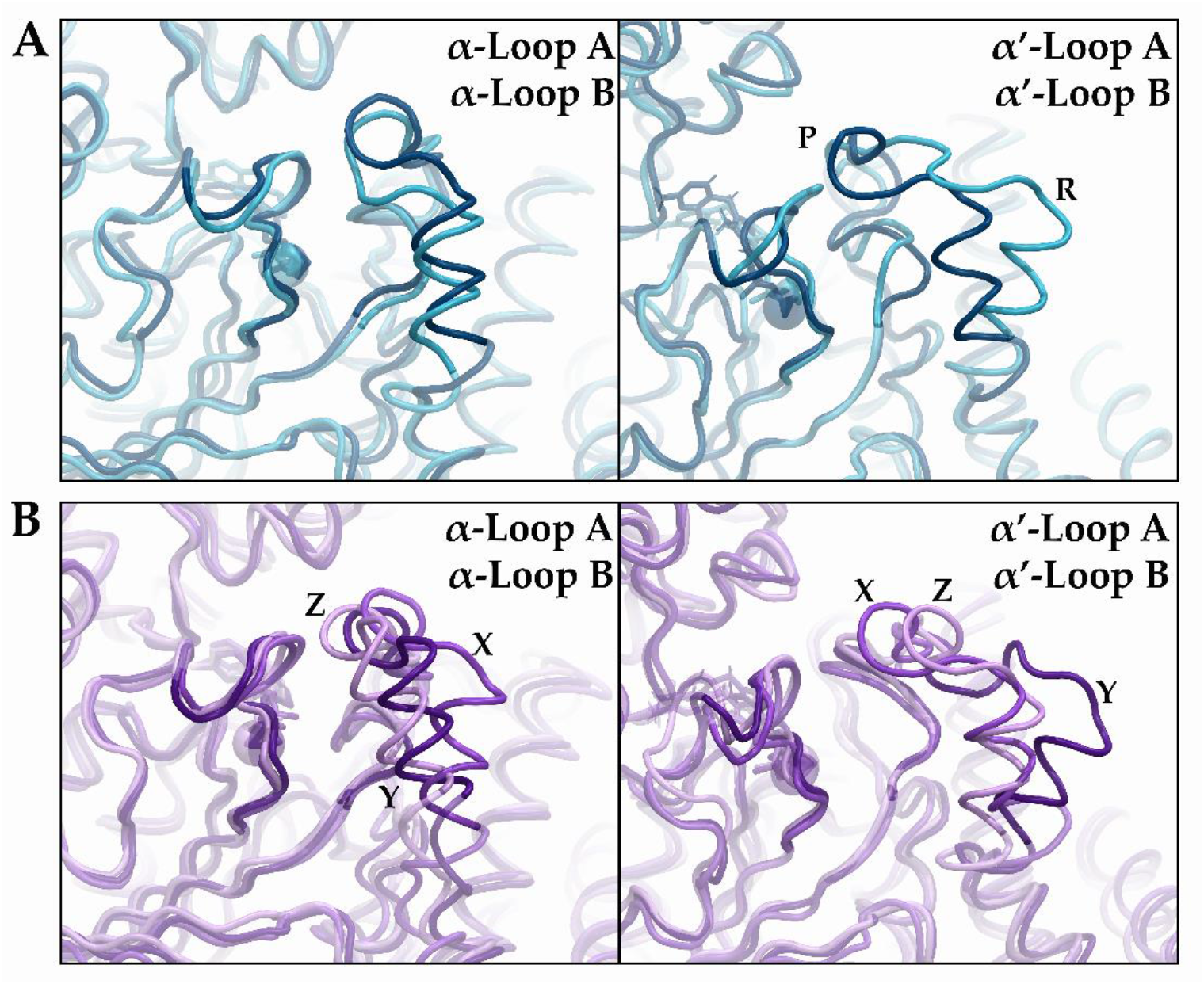
Clustering analysis indicates structures with different Loop A conformations on the α and α’ subunits. (A) Representatives of most populated (dark blue) and second most populated (light blue) clusters of wild-type (WT) E1 showing closed Loop A on the α subunit (left) and closed (P) or open (R) Loop A on the α’ subunit. (B) Representatives of most populated (dark purple, Y), second most populated (medium purple, X), and third most populated (light purple, Z) clusters of E1 variant with R349C replacement on both the α and α’ subunits showing closed or open Loop A on the α and α’ subunits. The locations of depicted representatives can be found on the density plots of principal component analysis in Figure 3A.

For E1 variants with DMMs, even though opening/closing actions of Loop A were observed in many of them, the motions of loops A and B change with the type of DMM (Figure 3 and Supplementary Figures S7-S20). For example, in contrast to WT E1, both loops (A and B) on the α and α’ subunits are actively involved in essential motions with R349C replacement on both the α and α’ subunits (Figure 3B). The behavior of R349C replacement on both the α and α’ subunits generates three different densely populated clusters, with conformational states corresponding to open or close loops (states X or Y, respectively), and other close loops on both subunits (state Z) (Figures 4B left and right panels). Although the direction of motion of the α’-Loop A with R349C replacement at the α and α’ subunits was similar to that of WT E1, the direction of motion of α-Loop A was almost orthogonal compared to WT E1 (Figure 3B).

In the case of E1 variant with R349H replacements on both the α and α’ subunits, PCA revealed that α-Loop A has a similar directional motion while α’-Loop A almost does not contribute to the essential motions (Figures S15 and S16). Clustering analysis of E1 variant with R349H replacements on both the α and α’ subunits (Supplementary Figure S16B) showed that the most populated clusters were conformationally similar to the state X (open α-Loop A and closed α’-Loop A) of E1 variant with R349C replacements on both the α and α’ subunits (Figure 4B). Interestingly, similar average reduction in cultured fibroblast-based PDC activity in males affected with PDCD is observed with R349C (21.5% control mean activity, min/max 8/40, STD 13.6, n=5)^30,59^ and R349H (21.2% control mean activity, min/max 10/50, STD 11.7, n=11)^30,59^. These results in conjunction with similar average reductions in PDC activity (for R349C and R349H replacements in male) indicate that Loop A may not only be important for anchoring the ligand (and hence the enzymatic catalysis), but may also be important for substrate intake and/or acetyl transfer to the E2. Since only the motion of Loop A on the α subunit resembles each other for these variants, a similar reduction of activity may stem from Loop A dynamics that affects one or a combination of three PDCD processes involving E1 (substrate intake to the active site of E1, reaction catalysis in the active site of E1, and/or transfer of acetyl from the active site of E1 to E2).

Unlike R349C replacements in the α and α’ subunits, PCA of E1 variant with R349C replacement at the α subunit only showed that even though the motion of the α-Loop A is similar to the WT E1, it demonstrated a perpendicular motion of α’-Loop A compared to WT (Figures S7 and S8). Cluster analysis of this E1 variant revealed that the closed conformational state was densely populated and that the open conformational state was almost nonexistent (Supplementary Figure S8). Interestingly, PDC activity from a single female cultured fibroblast specimen with R349C replacement was 15.3% control mean, but it remains to be determined what the variability in PDC activity in females might be after analysis of more specimens with R349C replacement from different individuals.

For E1 variant with W185C replacement in the α subunit only, the essential motions of Loop A on both the α and α’ subunits changed relative to WT E1 (Supplementary Figure S11A and B), and the close conformational state was highly populated (Supplementary Figure S12A). However, the disordering of the loops generated smaller clusters which also triggered conformational changes of TPP, especially in the α’ subunit (Supplementary Figure S12B right panel, most populated cluster representative colored with red). Interestingly, conformational changes in TPP are also observed with cluster analysis of W185C replacements on both the α and α’ subunits (Supplementary Figure S18B left panel, second most populated cluster representative colored with yellow).

Changes in the motions of Loops A and B can impact PDC-E1 activity. First, the cofactor TPP may not be held in a proper conformation in the active site. Examples of this are observed in some cluster representatives with replacements of W185C on one or both α subunits (Supplementary Figures S12 and S18, respectively). Second, the change in the direction of motions, such as with R349H replacement on the α subunit only (Supplementary Figure S9), could reduce the catalytic activity of E1 via disruptions of the active site–cofactor–substrate interactions or result in conformational changes of substrate or intermediate. Third, loop motions promote a closed active site channel with R349C replacement on the α subunit only (Supplementary Figure S8) or result in a channel with an improper conformational state. In that case, the lipoyl domain may not have access to the active site. In such instances, pyruvate metabolism might be interrupted or ineffective at the active site of E1, with a consequential reduction of PDC activity.

### 3.3 Dynamic Cross Correlation and the Centrality Analysis

Dynamic cross-correlation (DCC) analysis (Figure 5) provided information on highly correlated regions and the effect of DMMs on E1 dynamics. DCC of WT E1 revealed that multiple regions’ motion on different subunits correlated with each other (Figure 5A). High correlations were observed for regions involving amino acid stretches R29-G68 (Correlation Region 1, CR1), D286-A301 (CR2), and S302-D327 (CR3), all on both the α and α’ subunits (Figure 5B). A similar pattern was observed for all E1 variants with DMMs (Supplementary Figures S21-S24). CR1 is in proximity to the active site and packed in between CR2 and CR3, where Loop A is proximal to the region of CR2 (Figure 5B). CR1 contains residues within SSIC such as E1α R59, which interacts with E1β’ E309.^59^ In WT E1 and in almost all E1α variants with DMMs, the hydrogen bond interaction between E1α R59 and E1β’ E309 was conserved for about 90% of simulation time (data not shown). It is noteworthy that mutations of E1α R59 cause a reduction of PDC activity in both females (R59C, PDC activity is 40% of control mean)^66^ and males (R59S, PDC activity is 10% of control mean)^22^. E1α H63 within CR1 interacts with a substrate analog and is proposed to have a role in the reaction mechanism and E1α H63D or H63N replacements result in fibroblast-based PDC activity of ≤1% control mean.^41^ The change in the correlation between CR1, CR2, and CR3 as well as their correlation with other subunits in E1 variants with DMMs, suggest changes in E1 dynamics. For comparison, relative correlations for each E1 variant (Δ*Correlation* = *Variant^DCC^ – WT^DCC^*) were computed (Figure 5C and Supplementary Figures S23 and S24). We observed that DMMs altered the correlations of CR1, CR2, and CR3 indicating that both E1α replacements of R349 and W185 allosterically affect both the E1 active site and regions in its proximity.

**Figure 5.**
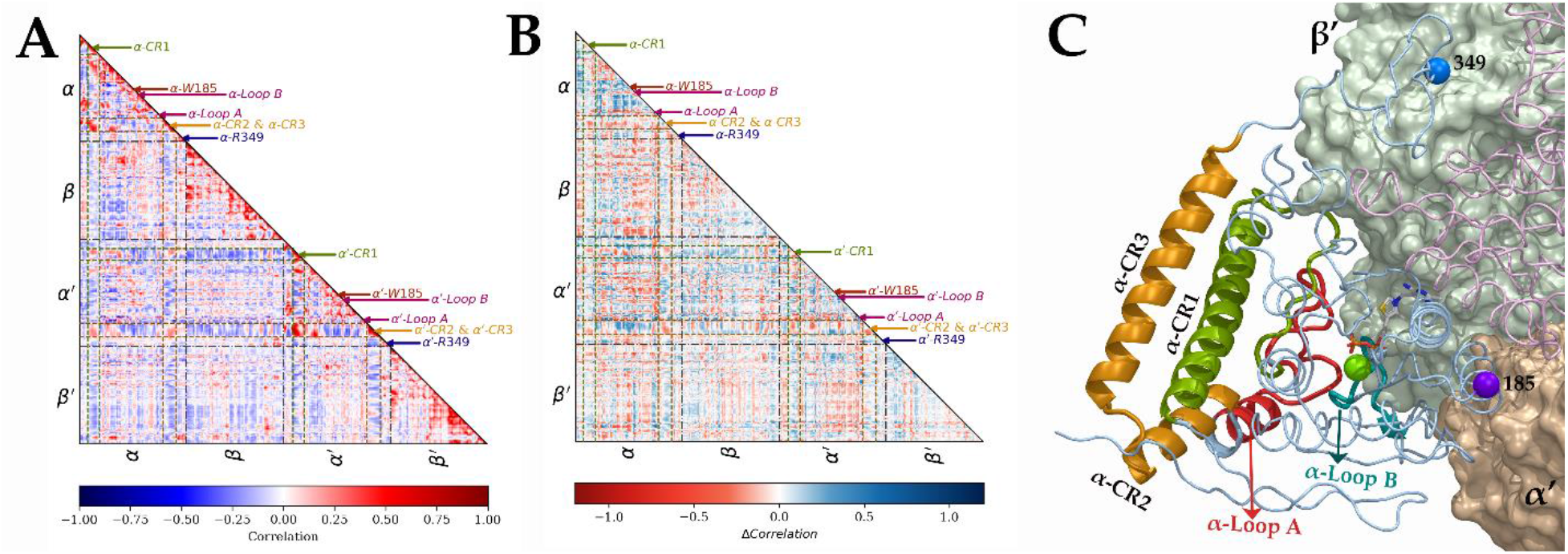
Lower triangular dynamic cross-correlation matrices of (A) wild type (WT) E1 and (B) E1 variant with W185R replacement on the α subunit alone relative to WT (Δ*Correlation = Variant_correlation_ – WT_correlation_*). Locations of the phosphorylation loops, the mutation regions, and the helices in close proximity to Loop A on the matrix are depicted with arrows. (C) Three-dimensional representation of part of WT E1 showing correlation region 1 (CR1, green; R29-Q68), correlation regions 2 and 3 (orange; CR2 and CR3 are consisting of D286-A301and S302-D327, respectively), Loop A (red), Loop B (cyan), W185 (sphere, purple), and R349 (sphere, blue).

Betweenness and eigenvector centrality calculations were performed to identify residues that influence the information flow and are essential for the communication between the amino acids of E1 (Figure 6). The effect of specific DMMs on the information flow was investigated using the relative centrality values (Δ*Centrality* = *Variant^C^ – WT^C^*, Supplementary Figures S25-S28). We observed that many residues located at SSIC have higher betweenness centrality (BC) and eigenvector centrality (EC) values relative to residues not involved in SSIC in WT E1.

**Figure 6.**
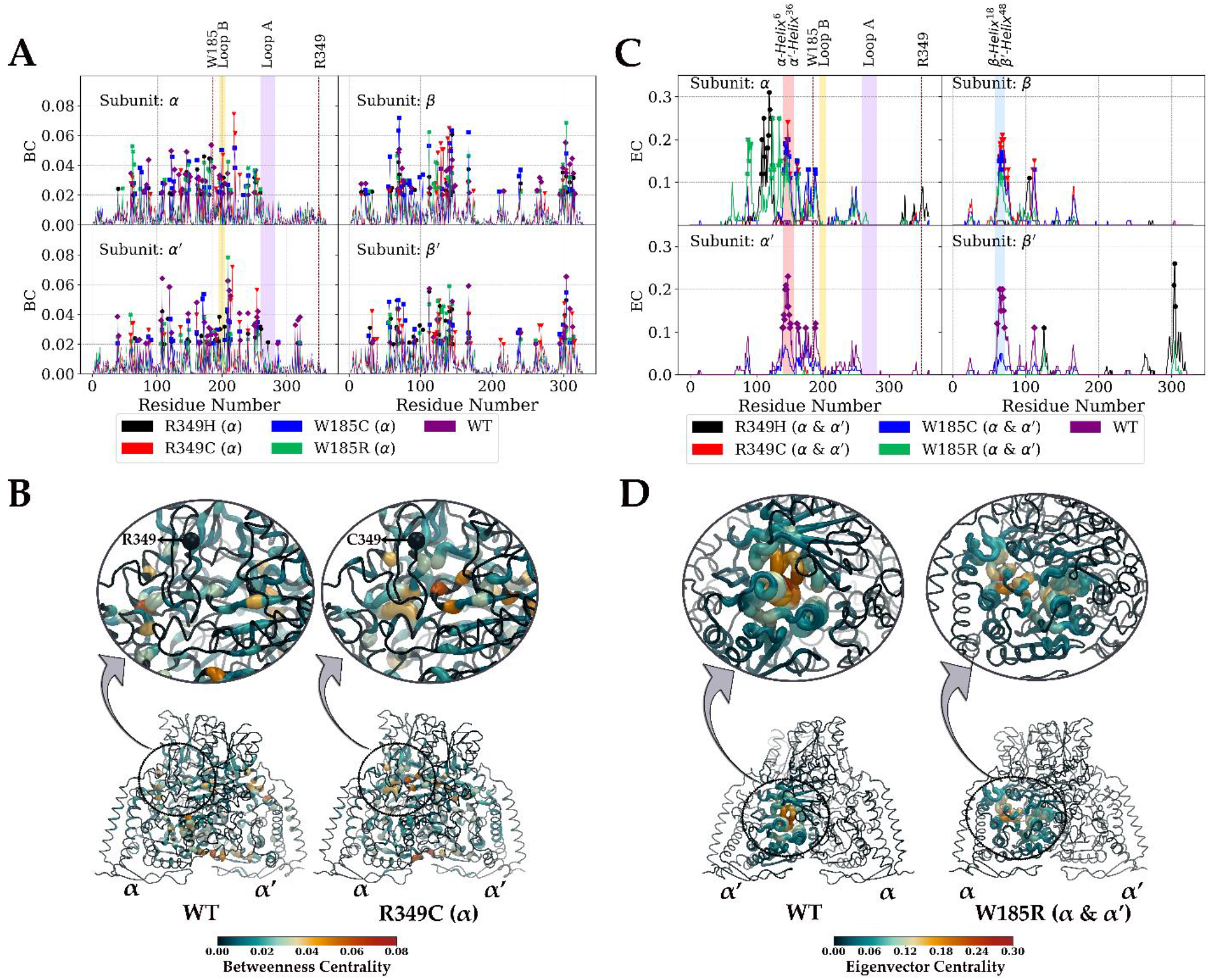
Centrality values show amino acid communication networks in wild-type (WT) E1 and the alterations to these networks with missense mutations. (A) Betweenness centrality values of WT E1 and E1 variants with DMMs on only α subunit. *BC* ≥ 0.02 are depicted with point markers. (B) Betweenness centrality values mapped on average simulations structures of WT E1 (left) and E1 variant with R349C replacement on only the α subunit (right). (C) Eigenvalue centrality values of WT E1 and E1 variants with DMMs on both the α and α’ subunits. (D) Eigenvalue centrality values mapped on average simulations structures of WT E1 (left) and E1 variant with W185R replacement on the α and α’ subunits (right). *EC* ≥ 0.1 are depicted with point markers.

Although slight differences were noted between BC of E1 variants with different DMMs, they all shared a common element; namely that BC of multiple residues located at SSIC changed (loss or gain) in almost all E1 variants (Figures 6A and 6B, and Supplementary Figures S25 and S26). For instance, residues R59, F61, E108, G119, G121 on the α and α’ subunits and residues M302, P303, Y304, E309, S312, I313 on the β’ and β subunits are part of the same SSIC region. E1β’ Y304 is packed in between residues E1α R59, E108, G119, and G121, while E1α R59 side chain interacts with E1β’ E309 side chain. The BC values of these residues changed depending on the type of DMM. For example, the BC value of E1α R59 decreased in all E1 variants with replacements on the α subunit alone, particularly E1 variants with W185C and W185R replacements (−0.01 ≤ ΔBC ≤ −0.03, Supplementary Figure S25), while the BC value for E1α G121 (−0.03 ≤ Δ*BC* ≤ −0.05) and E1β’ Y304 (−0.01 ≤ Δ*BC* ≤ −0.05) decreased in all E1 variants (Supplementary Figures S25 and S26). It should be noted that E1α G121 is located in the SSIC region formed by α, β and β’ subunits, while E1β’ Y304 is packed in between E1α G121 and E1α R59. Since these three residues are packed together in the same region, their corresponding ΔBC values are most likely correlated with each other, while higher ΔBC in E1α G121 relative to E1β’ Y304 and E1α R59 likely stems from its location on the SSIC formed by three subunits. Residues V124-Q127 on the β and β’ subunits involved in SSIC formed by α, β and β’ subunits were also affected by DMMs, particularly with R349C replacement on the α subunit (Figure 6A and 6B, ΔBC ≥ 0.03). In WT E1, this specific region almost had no influence in the communication network, but with R349C replacement on the α subunit, they gained influence in the information passing.

Another SSIC region with altered BC compared to WT E1 involved G173, A183, A209, R216, and D218 on α and α’ subunits. In WT E1, A183 and A209 on the α and α’ subunits, respectively, had high BC (BC = 0.05 and BC = 0.06, respectively), but in almost all E1 variants with DMMs, they lose their BCs (−0.01 ≤ ΔBC ≤ −0.05 A183 on the α subunit and −0.04 ≤ ΔBC ≤ −0.06 for A209 on the α’ subunit). In contrast, R216 and D218 on the α’ and α subunits had almost no BC in WT E1. Nevertheless, in the E1 variant with R349C replacement on the α subunit, these same residues were involved in information passing with a BC gain of 0.07.

In WT E1, residues A140-A151 (proximal end of α-Helix 6) on the α’ subunit and E59-A72 (α-Helix 3) on the β’ subunit are regions that have high EC values (Figure 6C and 6D). These residues are located on a pair of interacting helices located on the α’β’ heterodimer SSIC and are proposed to have a critical role in the coordination of the active sites via flip-flop action in E1.^16^ These helices are packed with GΦXXG motif, which exists in many TPP-dependent enzymes.^61–76^ Interestingly, the DMMs on E1 triggered changes in the communication network formed by this region and its symmetry-related counterparts on αβ heterodimer SSIC (Supplementary Figures S27 and S28). Among almost all E1 variants with DMMs on both α and α’ subunits, the EC of A141-A151 on the α subunit gained influence (ΔBC ≈ 0.2) on information flow while the EC of A141-A151 on the α’ subunit lost influence (ΔBC ≈ −0.2) (Figure 6C and 6D, and Supplementary Figure S28). The same pattern was observed for the E1 variant with R349C replacement on the α subunit only (Supplementary Figure S27).

The α-Helix 3 formed by E59-A72 on the β’ subunit and its symmetry-related counterpart E59-A72 on the β subunit were also affected by the DMMs. With R349C replacement on the α subunit only, the α-Helix 3 (especially G64-A70) on the β’ subunit loses its influence on the communication network (Δ*EC* ≈ –0.15) while EC increased (Δ*EC* ≈ 0.15) on the β subunit. Similarly, with R349C replacement on both the α and α’ subunits, the α-Helix 3 on β’ subunit lost its influence (Δ*EC* ≥ –0.15) as with this replacement on one subunit, while the EC increased (Δ*EC* ≥ 0.15) on the β subunit (Supplementary Figure S28). Males with R349C replacement exhibit severe clinical phenotype of neonatal lactic acidosis, developmental delay and Leigh disease with fibroblast-based PDC activity of 21.5% control mean activity (min/max 8/40, STD 13.6, n=5).^30,59^ Females with R349H and R349C replacements also have severe neonatal/infantile phenotype as males with fibroblast-based PDC activity ranging from 15-45% control mean (n=3).^30^ The communication network involving α-Helix 3 on the β and β’ subunits is most likely correlated with the flip-flop action that coordinates the active sites. Hence, any alteration to the communication provided by this region can affect the flip-flop action. In conjunction with the reported clinical phenotypes, these results suggest that any mutation that alters the flip-flop action is most likely to have severe clinical consequences.

Similar results for EC values of α-Helix 3 on the β and β’ subunits were obtained with W185R and W185C replacements on both the α and α’ subunits (Figure 6C and 6D, and Supplementary Figure S28). In addition to α-Helix 3, the proximal residues A88-H92, G122-M126, and N130-N135 on the α subunit also gained influence in the communication network with W185R replacement on both the α and α’ subunits (Figure 6C and 6D, and Supplementary Figure S28). These regions are all in proximity to each other and A140-A151 (proximal end of α-Helix 6) on the α subunit. E1α amino acid stretch G122-M126 is located at the SSIC formed by α-β-β’, while amino acid stretch A88-H92 is situated on the active site of E1α. Notably, the side chains of E1α Y89 and R90 are anchors to the phosphate groups of TPP^16^, while E1α G136 and V138 are distal to amino acid stretch N130-N135. These residues interact with the pyridinium ring of TPP via their backbones.^16^ Males with E1α W185R replacement show fibroblast-based PDC activity of <50% control mean and clinical phenotype consisting of Leigh disease, severe developmental delay and lactic acidosis that can be temporarily controlled with high dose thiamine supplementation (200 mg/kg/day).^77,78^ Furthermore, females with the V138M replacement consistently show fibroblast-based PDC activity of <7% control mean.^79,80^ This PDC activity reduction is not due to reduction of the decarboxylation activity of E1 but from distortion of the phosphorylation loops and displacement of the diphosphate tail of TPP due to the mutant M138 side chain.^39^ Therefore, any change in the communication provided by residues in close proximity to or on the helices involved in flip-flop action would not only affect the coordination of the active site regions but also influence the catalysis in the active sites.

BC and EC results provide insight into the impact of E1α DMMs at SSIC. Disease-causing mutations do not only disrupt local SSIC (such as with W185R replacement on both α and α’ subunits). Still, they can also affect interface contacts at other SSIC (such as R349C replacement at the α subunit alone). The influence on the communication network of the helices proposed to be essential for the flip-flop action can change with the type of mutation at distant sites. These results show that E1α DMMs influence the active sites allosterically by altering the information flow.

### 3.4 Hydrogen Bond Interactions at Heterotetramer Interface

Alternating sites reactivity is a common feature of TPP-dependent enzymes involving communication between the active sites in E1.^81–89^ This communication can be achieved with flip-flop action of human PDC-E1 and involves amino acids A140-N155 on the α and α’ subunits and E59-A72 on the β and β’ subunits.^16^ A molecular switch and proton wire mechanism for this communication were also suggested as an alternative to the flip-flop action mechanism.^20^ The proton wire mechanism links the cofactors on the two active sites via a channel formed by residues at the heterotetramer interface.^20^ The existence of a proton wire mechanism was shown experimentally for *E. coli*,^20,85,89^ but its existence in human PDC-E1 is unclear. In *E. coli*, the proton wire involves water molecules and acidic residues E235 and E237 on the α and α’ subunits^20,86^, while in *Bacillus stearothermophilus*, acidic residues E88, D180, and E183 on the α and α’ subunits and D91 on the β and β’ subunits play a role in this mechanism of action.^20,83^ In human PDC-E1, residues Q174 and E177 on the α and α’ subunits and E59, Q88, and D91 on the β and β’ subunits are the homologous sites for the residues forming the proton wire in *E. coli and B. stearothermophilus*. It should be noted that, unlike the residues that form the proposed proton wire in human PDC-E1, the proton wire in both *E. coli and B. stearothermophilus* are formed by residues that are all acidic in nature.

In human PDC-E1, the proposed proton wire is located on the heterotetramer interface that encompasses the residues involved in the flip-flop action and the helices within stretches Q172-K186 on the α and α’ subunits, and T82-N95 on β and β’ subunits. Residues Q174 and E177 on α and α’ subunits and Q88 and D91 on β and β’ subunits in this interface are in close proximity to one another. To investigate changes in interactions within the heterotetramer interface with DMMs, the fractions of hydrogen bond formation between these specific residues were calculated along the trajectories of all the E1 variants (Figure 7).

**Figure 7.**
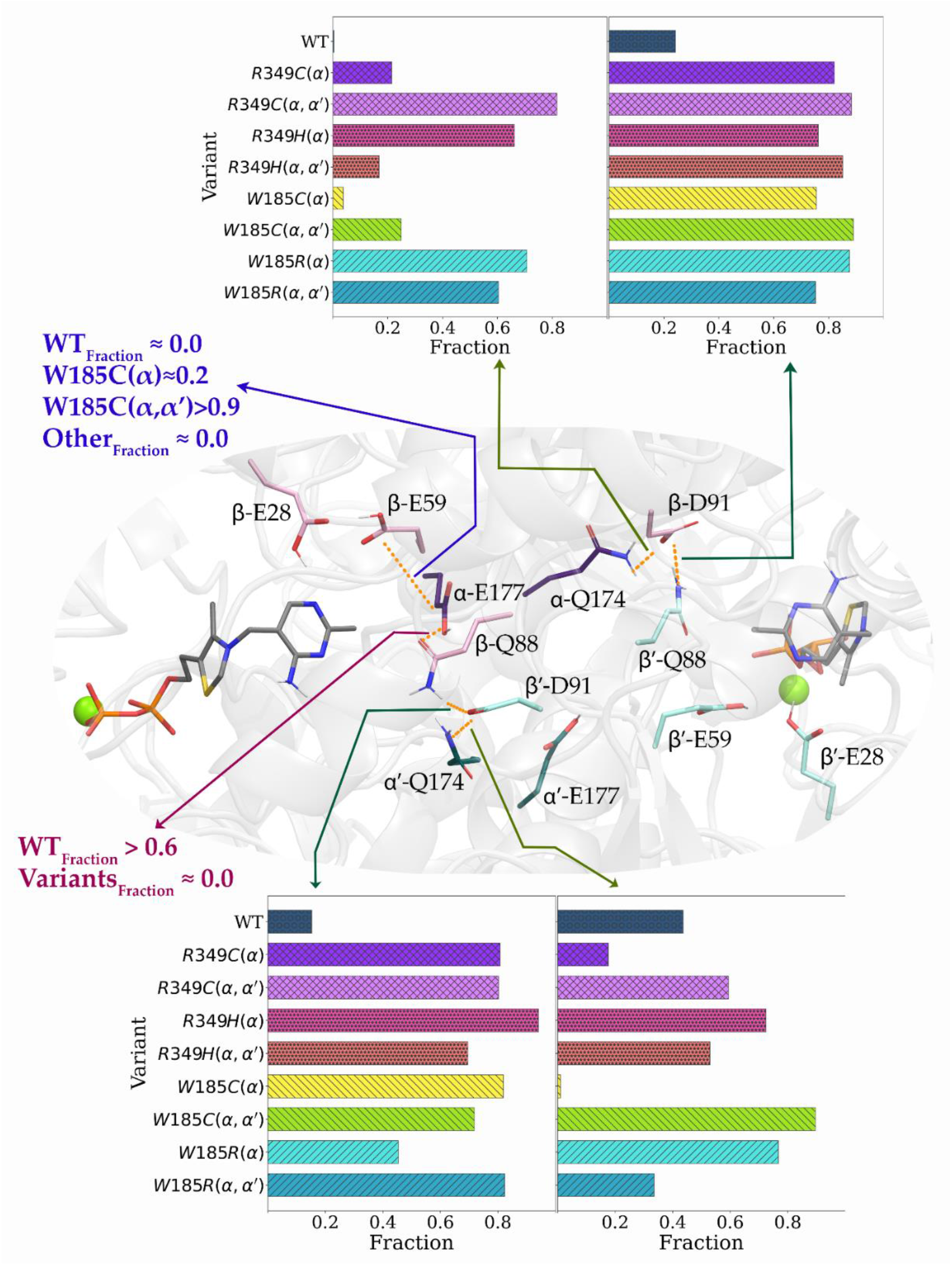
During the simulations, the fractions of hydrogen bond formation between the subunit interfaces indicate an increase in subunit-subunit interactions between the two active sites for almost all E1 variants compared to wild-type E1.

DMMs promote a shift of interactions at this heterotetramer interface. In WT E1, hydrogen bond formation is observed between E1α E177 and E1β Q88. However, this interaction almost ceased to exist with all the E1 variants. E1β Q88 forms another hydrogen bond with E1β’ D91 in about 20% of the WT E1 trajectory (Figure 7, bottom-left). A similar result was observed with their symmetry-related counterparts (Figure 7, top-right). Hydrogen bond formation fractions between these residues increased in E1 variants regardless of the DMM site or type (Figure 7). Furthermore, E1β’ D91 forms another hydrogen bond with E1α’ Q174 in about 50% of the WT E1 trajectory (Figure 7, bottom-right), while no hydrogen bond formation was observed for their symmetry-related counterparts (E1β D91 and E1α Q174, Figure 7, top-left). Hydrogen bond interactions between these sites were altered in E1 variants depending on the location and type of DMM (Figure 7). For instance, with R349C replacements on both α and α’ subunits, the E1β D91 and E1α Q174 hydrogen bond interaction was more abundant than in WT E1 or E1 variant with R349C replacement at the α subunit alone. With replacements at W185, the hydrogen bond interactions between E1β’ D91/E1α’ Q174, E1β D91/E1α Q174, and E1β D91/E1β’ Q88 are conserved longer in W185C replacement at both α subunits compared to W185C replacement at the α subunit alone, but the opposite was observed with the W185R replacements (Figure 7). Furthermore, W185C replacements on one or both α subunits resulted in an additional novel hydrogen bond interaction between E1α E177 and E1β E59, which was not observed in WT E1 or other E1 variants tested here.

These observations suggest that the hydrogen bond interactions at this heterotetramer interface in E1 changes depending on the type and/or location of the E1α DMM. The E1β Q88 almost always forms a hydrogen bond interaction either with the α or β’ subunit in WT E1, while it interacts only with the β’ subunit in E1 variants with DMMs. The results also suggest that the type of replacement at the same residue has different effects on hydrogen bond interaction at this heterotetramer interface. These results indicate that nearby or far-off residue replacements (such as at W185 or R349, respectively) can disrupt interactions in the heterotetramer interface in human PDC-E1. If a proton-wire exists in human PDC-E1 that links two active sites for activation/inactivation of TPP, then changes in the interaction profile in this heterotetramer interface would disrupt this link.

## 4. CONCLUSIONS AND FUTURE DIRECTIONS

We found that mobility of phosphorylation Loop A changes with DMMs. PCA revealed that the essential motions for the biological activity of E1 change with DMMs, with loops A and B as the major contributors to the essential motions. Cluster analysis showed that the α’ loops are either close or open in WT E1 and that these states are achieved by the essential motions observed with PCA. Change in the highly populated clusters with different conformational states could differentially impact E1 function. First, enzyme affinity for the substrate may change. Second, reaction catalysis may be hindered because loops A and B are important for achieving specific cofactor-ligand conformation. Third, acetyl transfer to E2 may be negatively affected either by close loops or the loss of lipoyl domain recognition. This suggests that similar PDC activity reductions could be observed due to one or a combination of scenarios mentioned above. Variation of the loop structure and its dynamics appears to be the main reason responsible for the functional PDC deficiency even though the specific DMM is not within the loop region.

Dynamic cross-correlation analysis showed that the motions of residues located at SSICs are correlated with each other, especially at regions in proximity to the active site. Betweenness centrality analysis with these correlations indicated that the communication in WT E1 is achieved by residues located at SSICs. Eigenvector centrality analysis suggested that α-helices 6 and 18 on the α and β subunits, respectively (and their symmetry-related counterparts) are important for communication. These helices were suggested as regions that control the flip-flop action for alternating site activity.^16^ These results suggest that the alternating site activity is most likely achieved by the flip-flop action controlled by the motion of these helices. Communication networks provided by eigenvector centrality analysis were altered by DMMs even if they are not in this region, suggesting that DMMs could trigger a change in the flip-flop action of E1, leading to reduced enzymatic activity. Hydrogen bond analysis revealed that the subunit interaction at the core of E1 increases with DMMs. This is most likely correlated with change(s) of the flip-flop action due to DMMs because several of the residues in this SSIC are located on the helices that regulate the flip-flop action.

Collectively, these results suggest an allosteric effect in PDC-E1 function with implication for the overall PDC activity and thus disease from DMMs. DMMs at SSICs prompt a series of changes in E1, such as essential motions and communication within the enzyme. These changes are correlated with different loops A and B conformational states. Undesired conformational states of these loops could decrease PDC activity by reducing 1) enzyme affinity for pyruvate, 2) catalysis in the active site, 3) recognition of lipoyl domain, and 4) substrate channeling, or some combination of the above.

Future work will focus on finding blood-brain-barrier permeable small molecule candidates that would partially or fully restore PDC activity of E1 variants with E1α DMMs to WT activity levels. We will virtually screen for candidates that induce the desired loop conformations for enzymatic catalysis by restoring original PDC-E1 dynamics in WT. Subsequently, we will test the candidate small molecules for their capacity to partially or fully restore *in vitro* PDC activity from human cell extracts with specific DMMs before embarking on animal-model evaluations.

## Acknowledgment

This research was supported in part by NIH 2U54NS078059-09 RDCRN NAMDC Project 4 grant (to JKB). We thank Nicole Ducich for discussions. HG and OI acknowledge support from CHE-2041108. HG and OI also acknowledge the Extreme Science and Engineering Discovery Environment (XSEDE) award CHE200122, which is supported by NSF grant number ACI-1053575. This research is part of the Frontera computing project at the Texas Advanced Computing Center. Frontera is made possible by the National Science Foundation award OAC-1818253. HG and OI gratefully acknowledge the support and hardware donation from NVIDIA Corporation and express our special gratitude to Jonathan Lefman.

