## Supporting Information for "Simulations of Pathogenic E1α Variants: Allostery and Impact on Pyruvate Dehydrogenase Complex-E1 Structure and Function"

### FOR

#### 1 RMSD

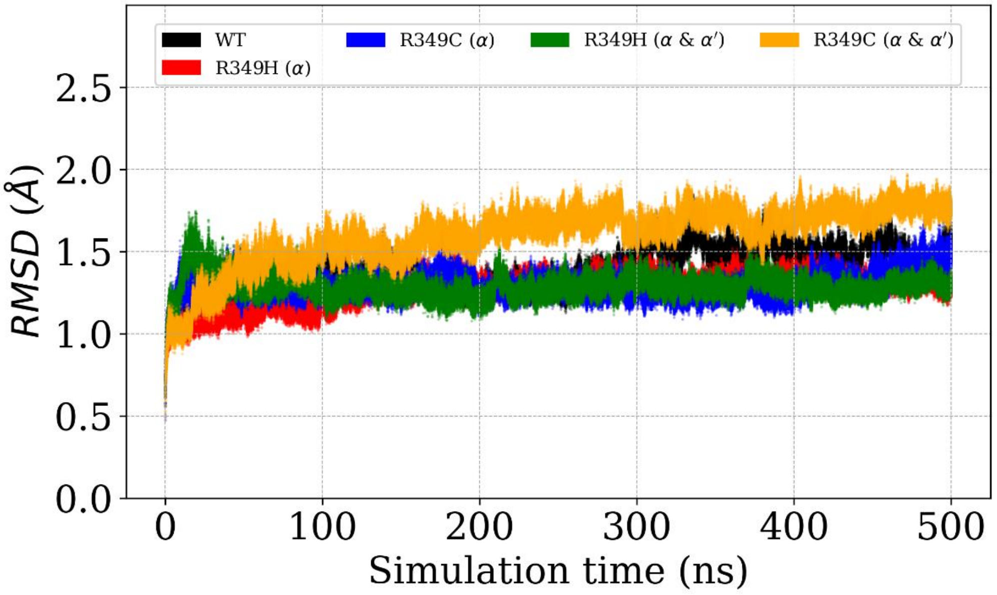

Figure S.1 Root mean square deviations (RMSD) of backbone atoms (C, $C^{\alpha}$, and N) along the simulation of WT, and variants with mutations on α subunit.

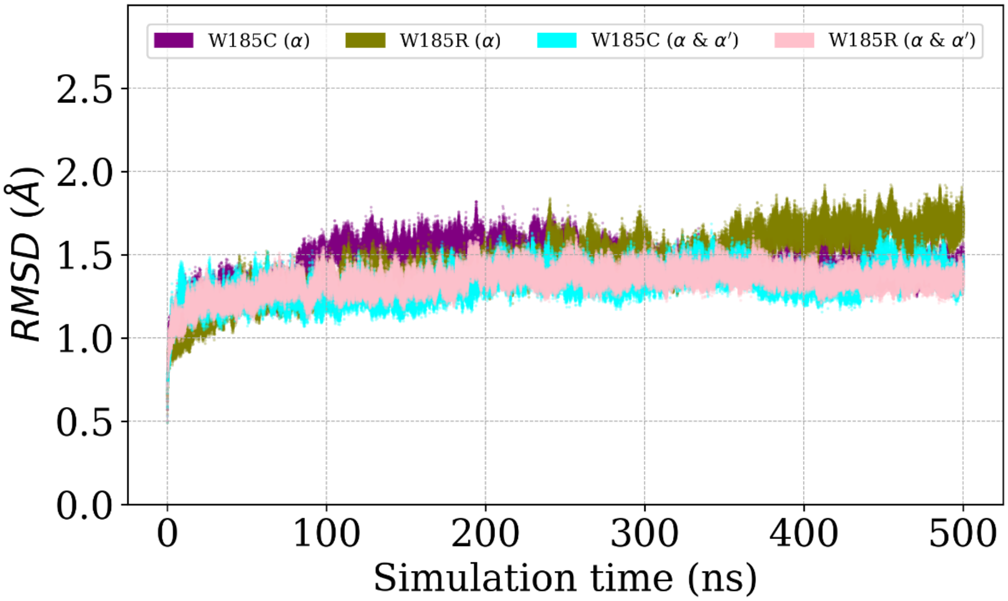

Figure S.2 Root mean square deviations (RMSD) of backbone atoms (C, $C^{\alpha}$, and N) along the simulations variants with mutations on α and α' subunits.

#### 2 RMSF

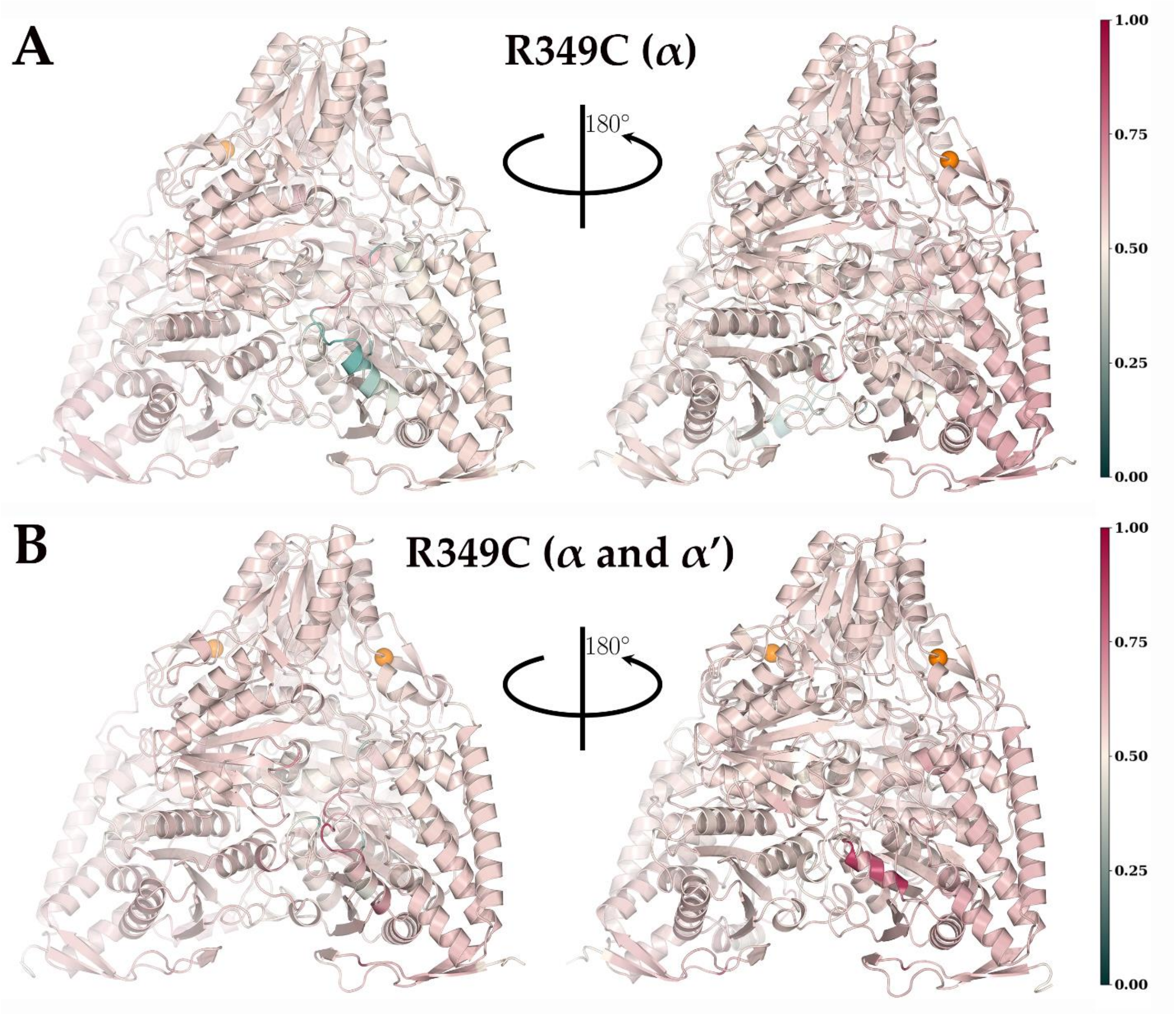

Figure S.3 Normalized relative RMSF values ($\Delta\widehat{RMSF} = \widehat{V_{rmsf} - WT_{rmsf}}$) of C$^\alpha$ atoms on (A) variant R349C (α) and (B) variant R349C (α and α’) shown on the three-dimensional representations of their corresponding average structures obtained from simulations. The locations of the mutations are represented with spheres.

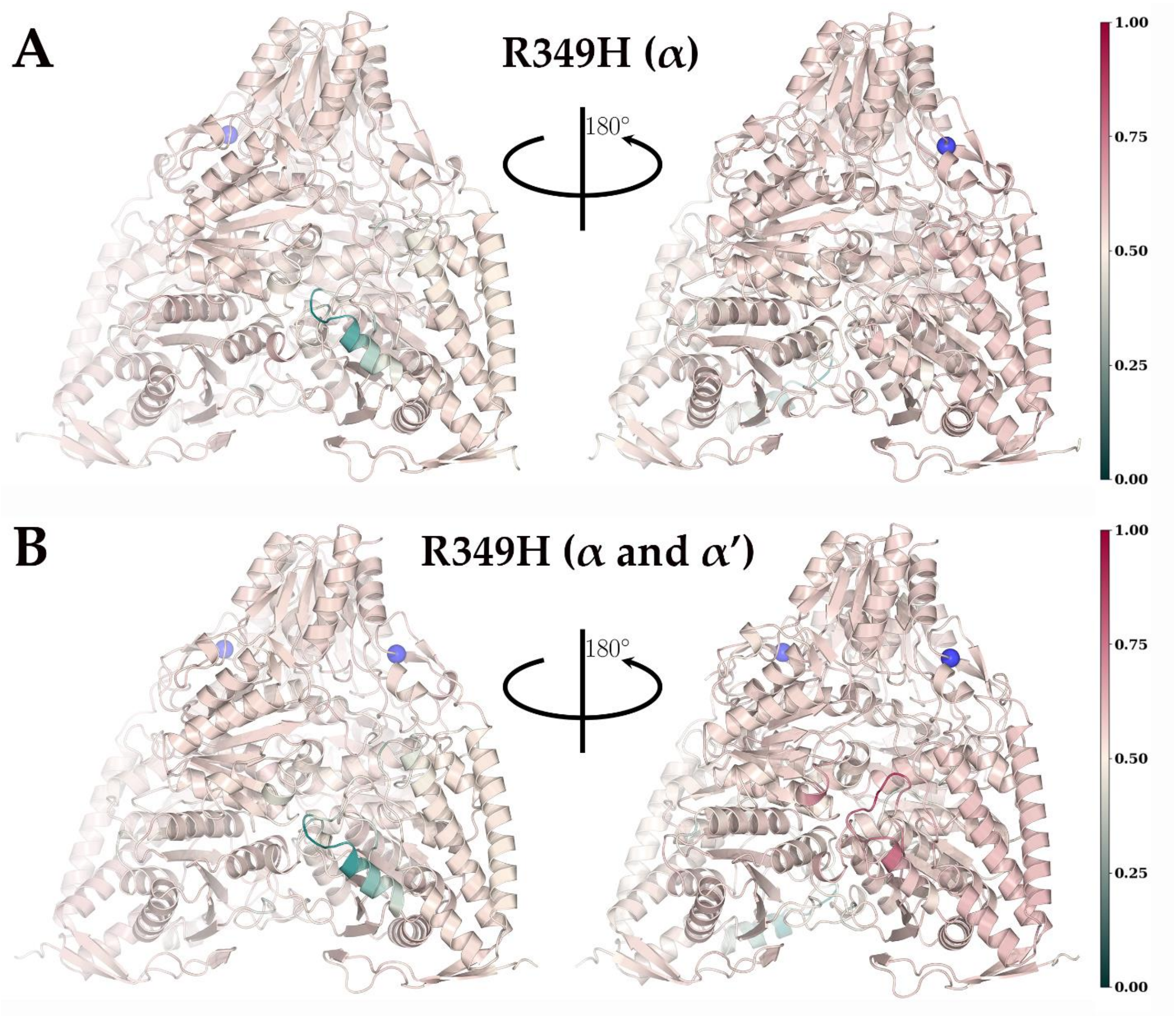

Figure S.4 Normalized relative RMSF values ($\Delta\widehat{RMSF} = V_{rmsf} \widehat{-WT}_{rmsf}$) of C$^{\alpha}$ atoms on (A) variant R349H (α) and (B) variant R349H (α and α’) shown on the three-dimensional representations of their corresponding average structures obtained from simulations. The locations of the mutations are represented with spheres.

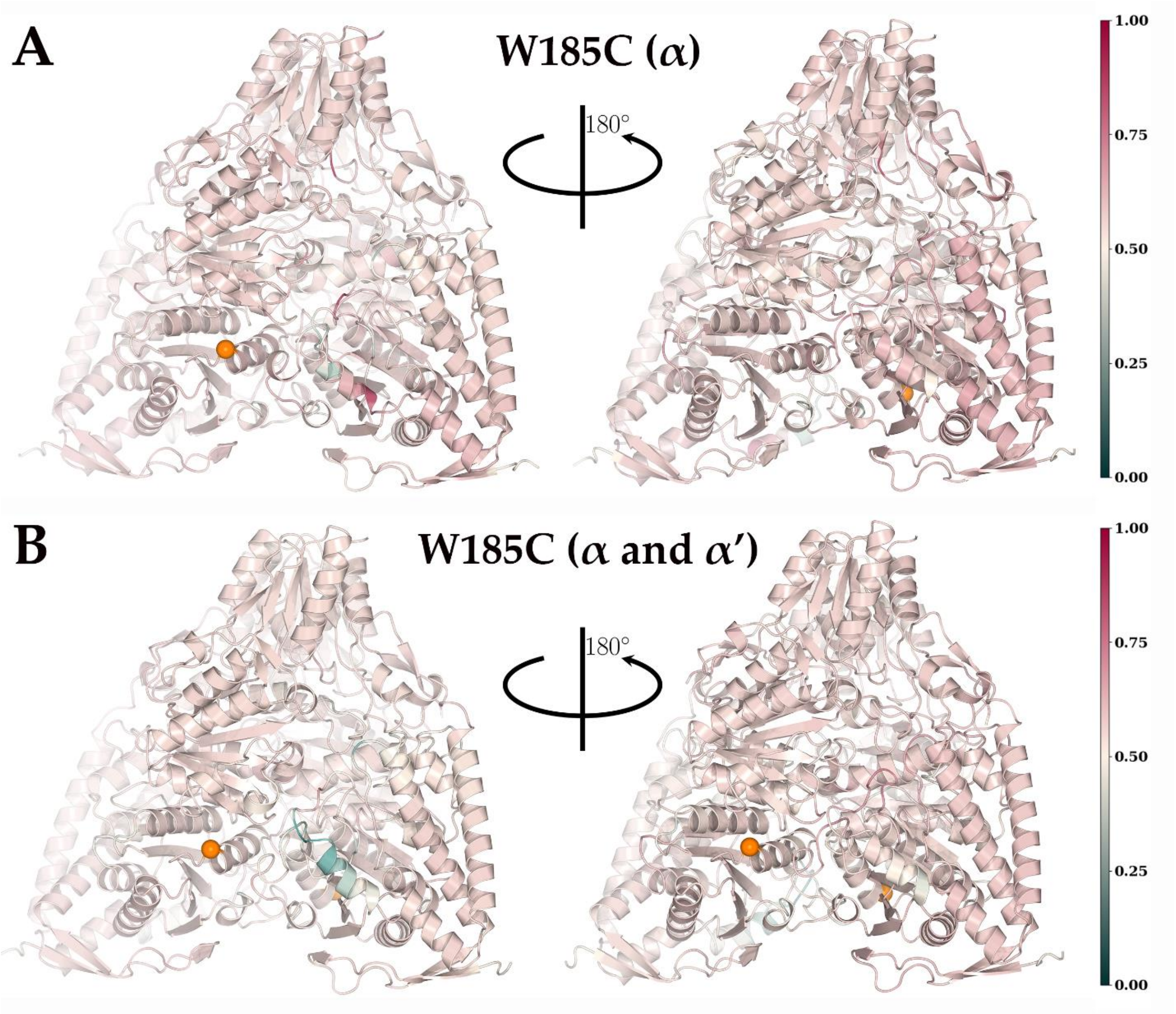

Figure S.5 Normalized relative RMSF values ($\Delta\widehat{RMSF} = V_{rmsf} \widehat{-WT}_{rmsf}$) of $C^{\alpha}$ atoms on (A) variant W185C (α) and (B) variant W185C (α and α') shown on the three-dimensional representations of their corresponding average structures obtained from simulations. The locations of the mutations are represented with spheres.

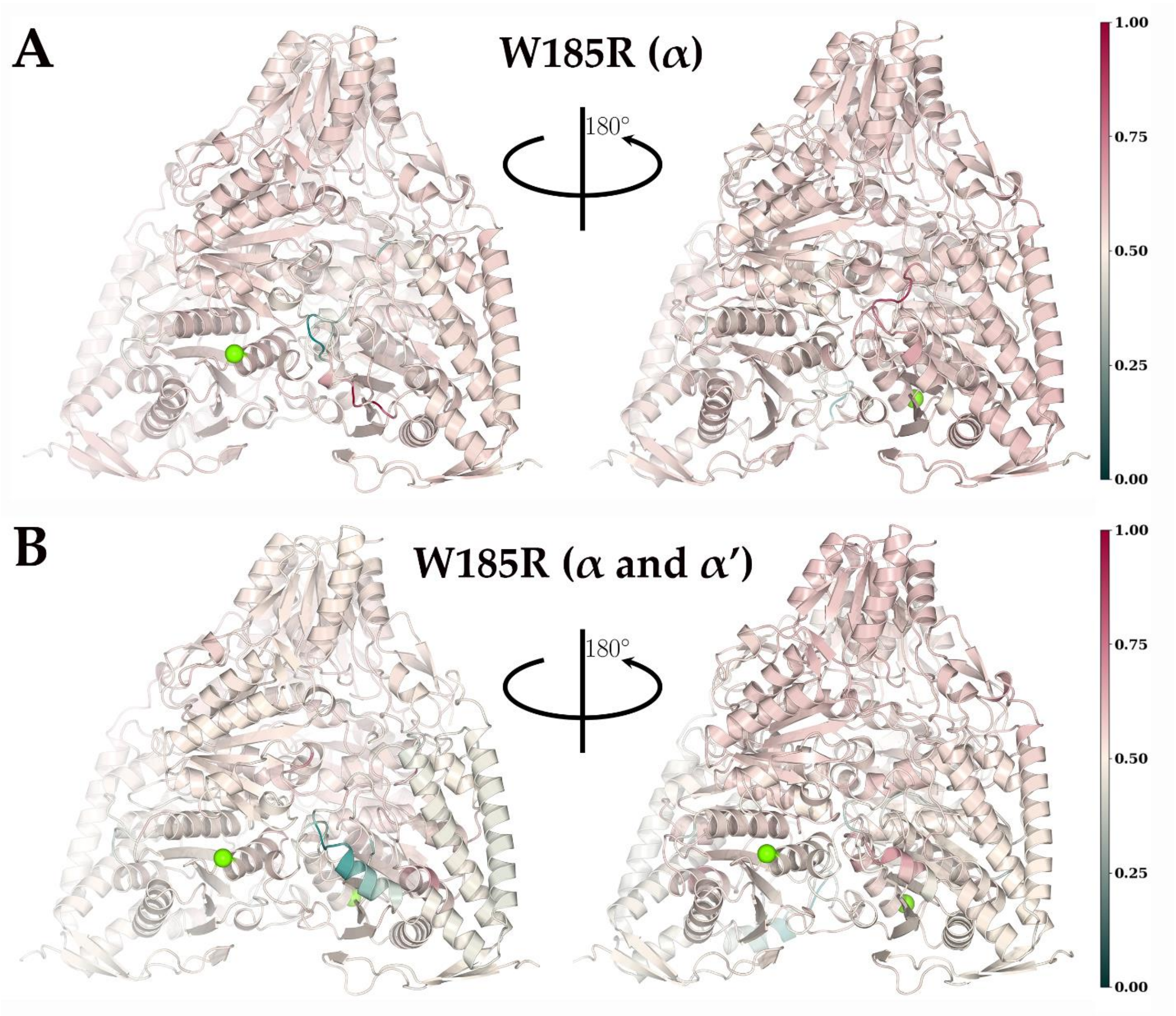

Figure S.6 Normalized relative RMSF values ($\Delta\widehat{RMSF} = V_{rmsf}\ \widehat{-WT}_{rmsf}$) of $C^{\alpha}$ atoms on (A) variant W185R (α) and (B) variant W185R (α and α') shown on the three-dimensional representations of their corresponding average structures obtained from simulations. The locations of the mutations are represented with spheres.

### 3 PRINCIPAL COMPONENT AND CLUSTER ANALYSIS

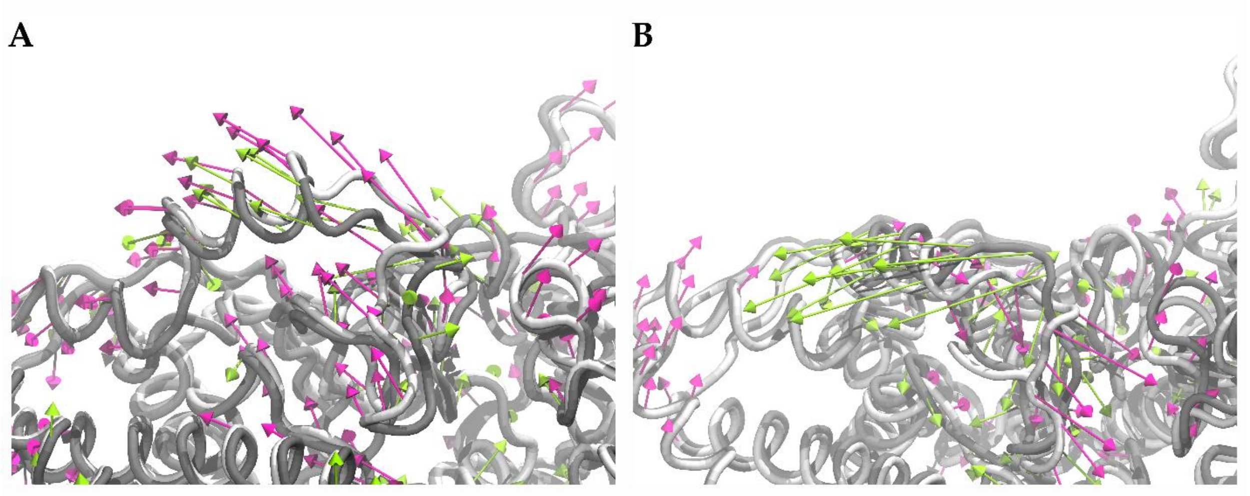

Figure S.7 First principal components (PC-1) of Loop A on A. α subunit, and B. α’ subunit showing alterations of R349C (α) variant (magenta) compared to WT E1 (green).

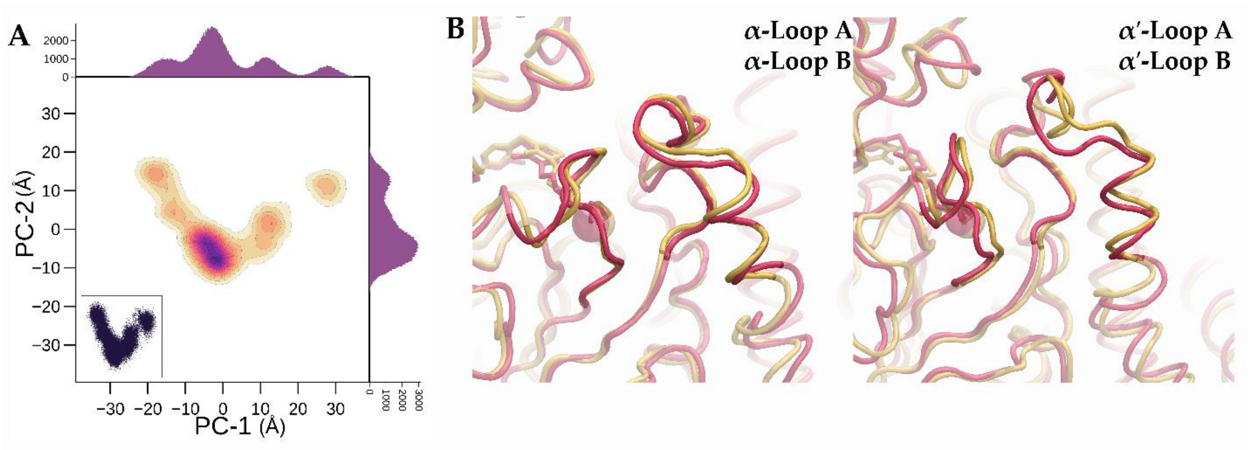

Figure S.8 Principal component analysis (PCA) and cluster analysis for R349C (α) variant. A. Two-dimensional density plots for the trajectory projected onto the subspace spanned by the first two principal components, PC-1 and PC-2. Scatter plot for same components is shown lower left corner while their populations are denoted with 2D histograms located at top (PC-1) and right (PC-2). B. Representative structures obtained from cluster analysis. Most populated cluster and the second most populated cluster representatives are depicted with red and yellow colors, respectively. The phosphorylation loops, Loop A and Loop B, located on α subunit is shown left and their counterparts on α’ subunit are depicted on right.

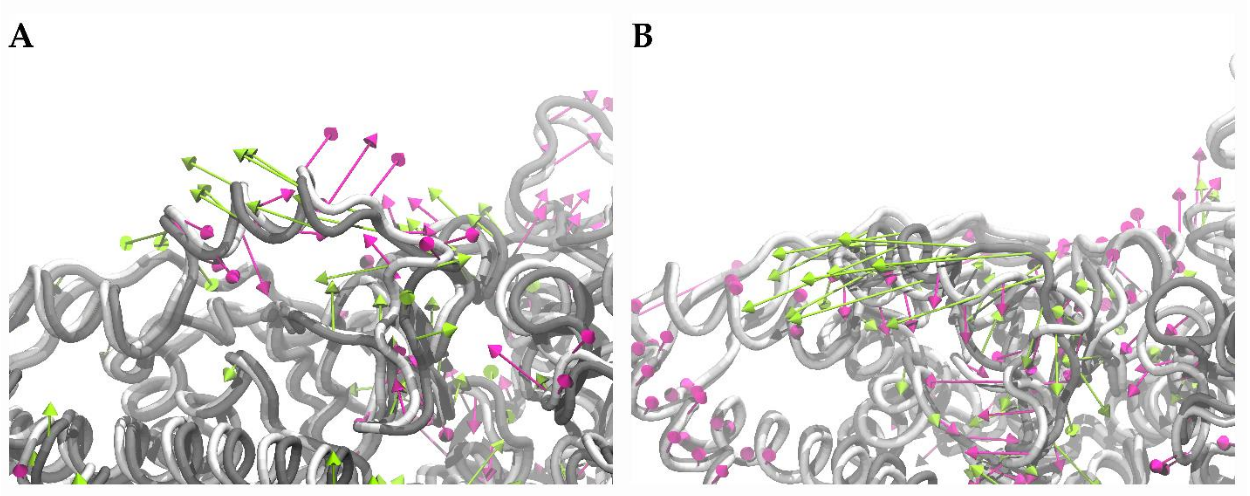

Figure S.9 First principal components (PC-1) of Loop A on A. α subunit, and B. α’ subunit showing alterations of R349H (α) variant (magenta) compared to WT E1 (green).

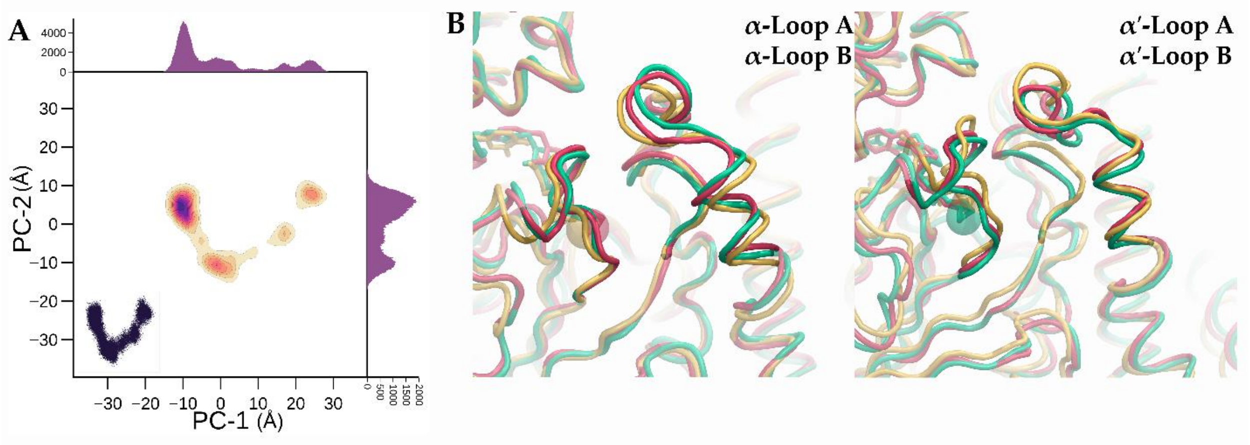

Figure S.10 Principal component analysis (PCA) and cluster analysis for R349H (α) variant. A. Two-dimensional density plots for the trajectory projected onto the subspace spanned by the first two principal components, PC-1 and PC-2. Scatter plot for same components is shown lower left corner while their populations are denoted with 2D histograms located at top (PC-1) and right (PC-2). B. Representative structures obtained from cluster analysis. Most populated cluster representative is shown with red while the second and third most populated clusters representatives are depicted with yellow and green colors, respectively. The phosphorylation loops, Loop A and Loop B, located on α subunit is shown left and their counterparts on α’ subunit are depicted on right.

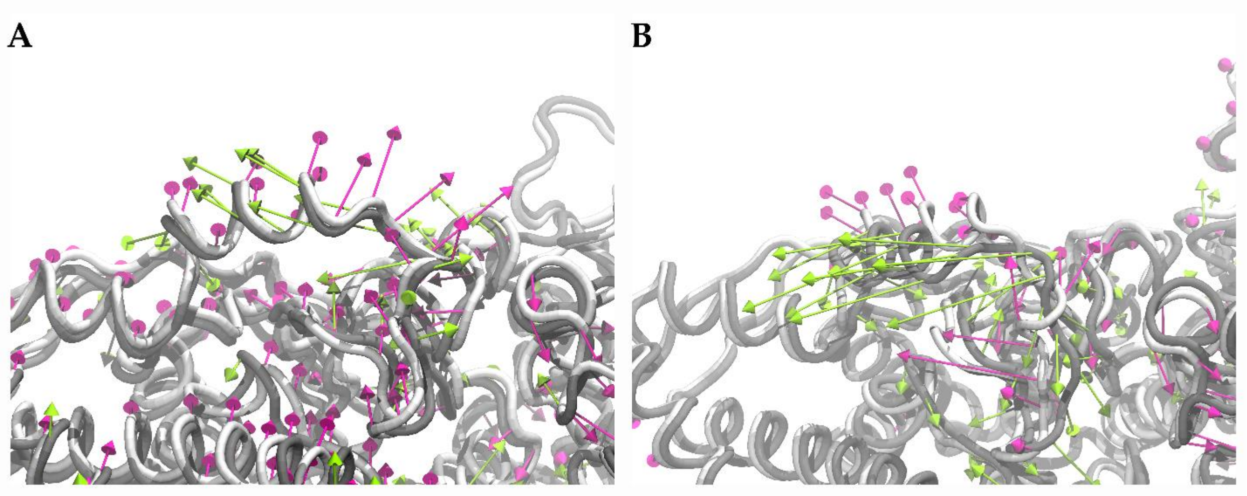

Figure S.11 First principal components (PC-1) of Loop A on A. α subunit, and B. α’ subunit showing alterations of W185C (α) variant (magenta) compared to WT E1 (green).

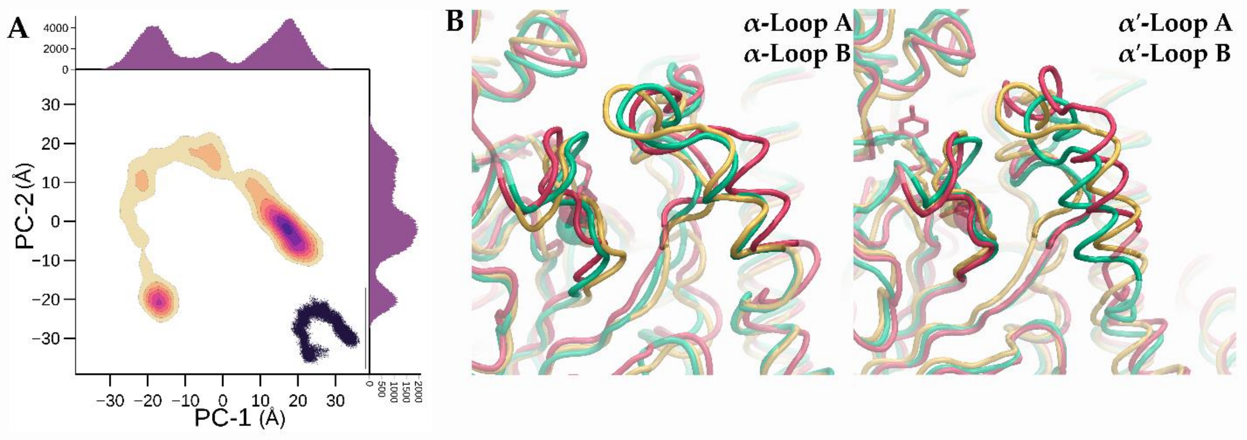

Figure S.12 Principal component analysis (PCA) and cluster analysis for W185C (α) variant. A. Two-dimensional density plots for the trajectory projected onto the subspace spanned by the first two principal components, PC-1 and PC-2. Scatter plot for same components is shown lower left corner while their populations are denoted with 2D histograms located at top (PC-1) and right (PC-2). B. Representative structures obtained from cluster analysis. Most populated cluster representative is shown with red while the second and third most populated clusters representatives are depicted with yellow and green colors, respectively. The phosphorylation loops, Loop A and Loop B, located on α subunit is shown left and their counterparts on α’ subunit are depicted on right.

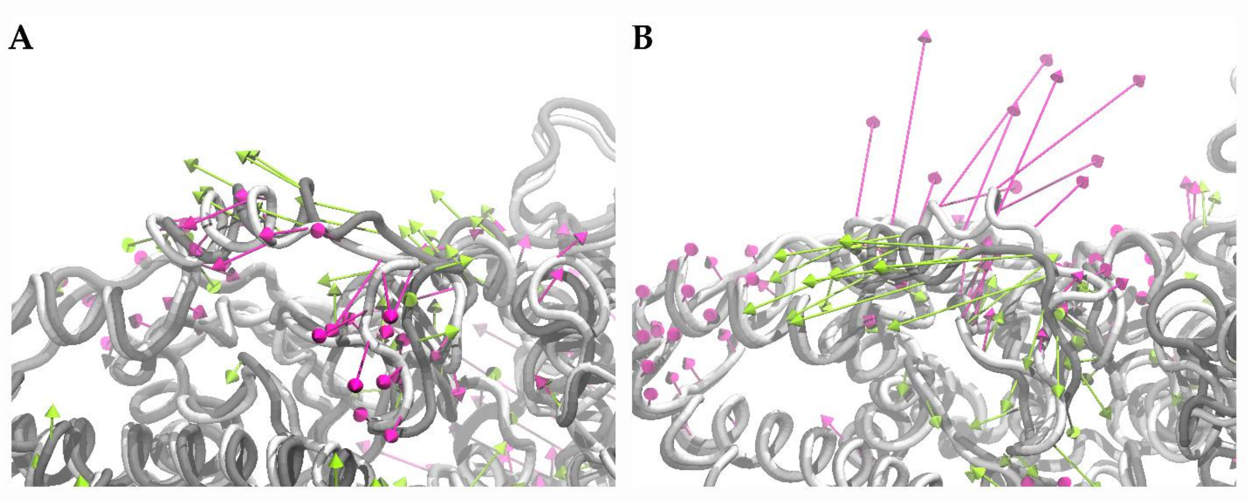

Figure S13 First principal components (PC-1) of Loop A on A. α subunit, and B. α’ subunit showing alterations of W185R (α) variant (magenta) compared to WT E1 (green).

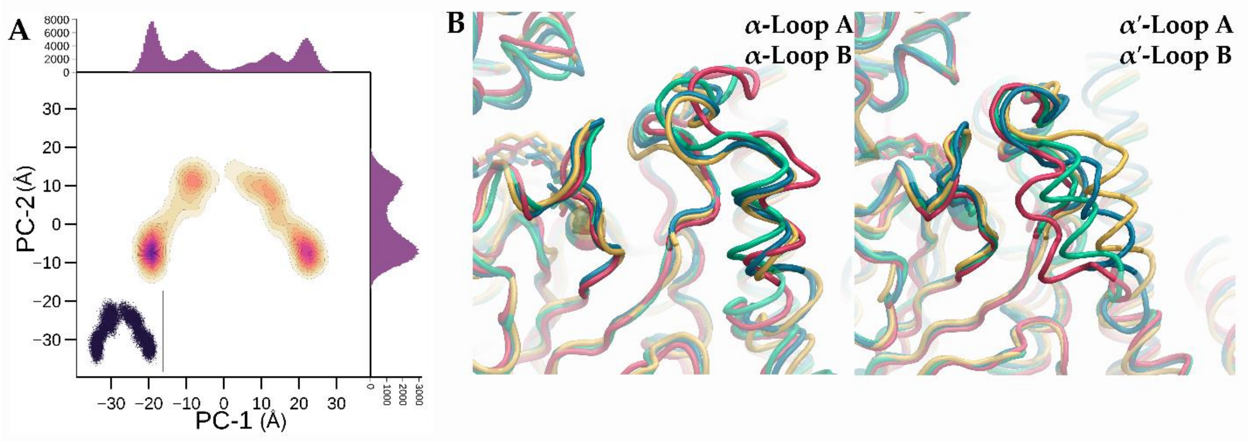

Figure S14 Principal component analysis (PCA) and cluster analysis for W185R (α) variant. A. Two-dimensional density plots for the trajectory projected onto the subspace spanned by the first two principal components, PC-1 and PC-2. Scatter plot for same components is shown lower left corner while their populations are denoted with 2D histograms located at top (PC-1) and right (PC-2). B. Representative structures obtained from cluster analysis. Most populated cluster representative is shown with red while the second, the third and the fourth most populated clusters representatives are depicted with yellow, green and blue colors, respectively. The phosphorylation loops, Loop A and Loop B, located on α subunit is shown left and their counterparts on α’ subunit are depicted on right.

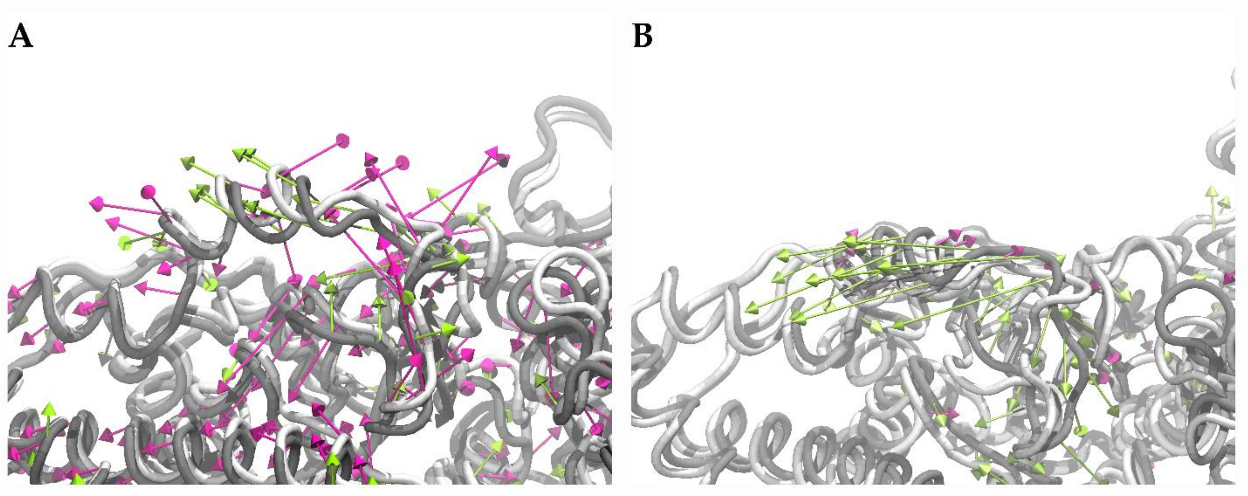

Figure S15 First principal components (PC-1) of Loop A on A. α subunit, and B. α' subunit showing alterations of R349H (α & α') variant (magenta) compared to WT E1 (green).

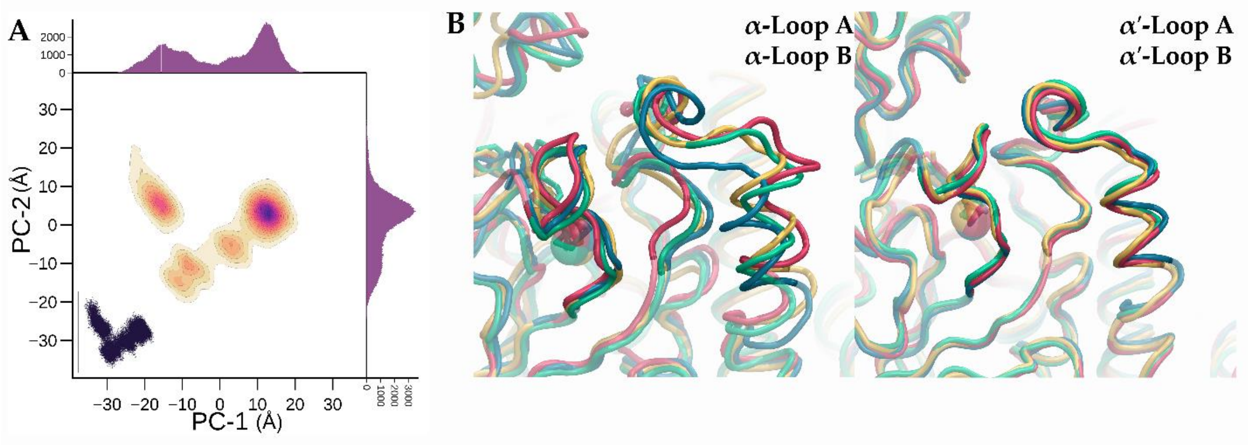

Figure S16 Principal component analysis (PCA) and cluster analysis for R349H (α & α') variant. A. Two-dimensional density plots for the trajectory projected onto the subspace spanned by the first two principal components, PC-1 and PC-2. Scatter plot for same components is shown lower left corner while their populations are denoted with 2D histograms located at top (PC-1) and right (PC-2). B. Representative structures obtained from cluster analysis. Most populated cluster representative is shown with red while the second, the third and the fourth most populated clusters representatives are depicted with yellow, green and blue colors, respectively. The phosphorylation loops, Loop A and Loop B, located on α subunit is shown left and their counterparts on α' subunit are depicted on right.

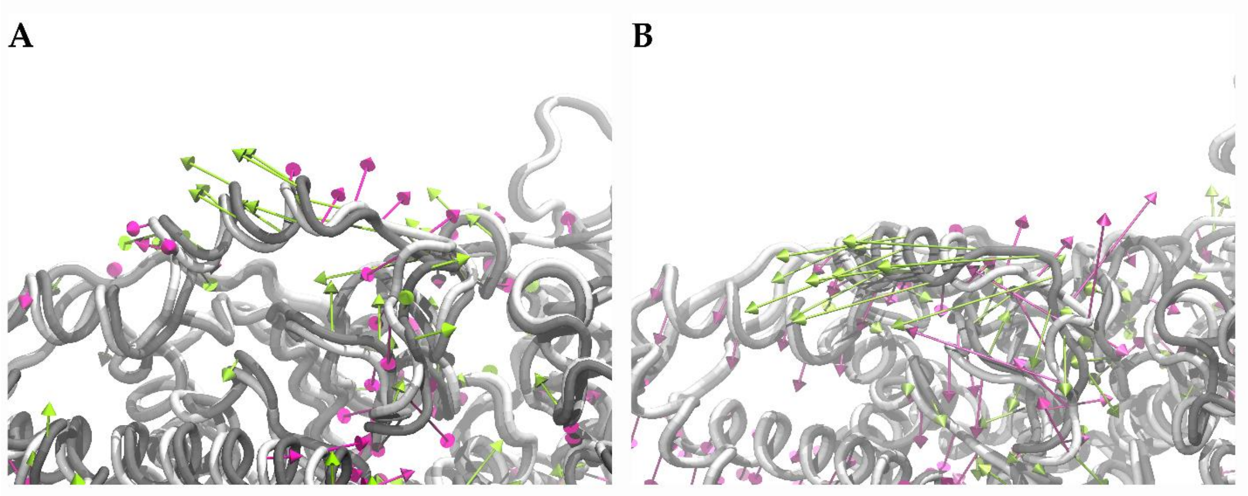

Figure S17 First principal components (PC-1) of Loop A on A. α subunit, and B. α' subunit showing alterations of W185C (α & α') variant (magenta) compared to WT E1 (green).

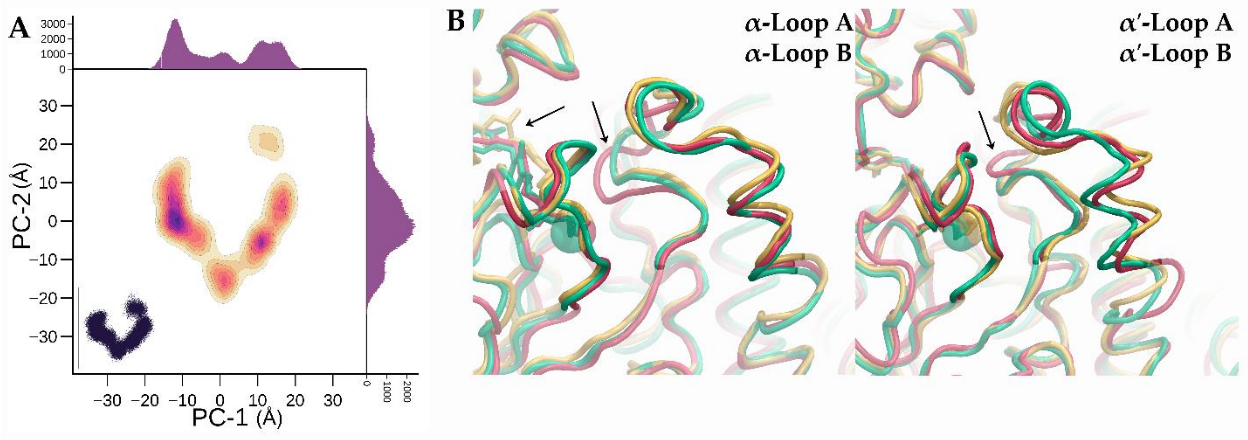

Figure S18 Principal component analysis (PCA) and cluster analysis for W185C (α & α') variant. A. Two-dimensional density plots for the trajectory projected onto the subspace spanned by the first two principal components, PC-1 and PC-2. Scatter plot for same components is shown lower left corner while their populations are denoted with 2D histograms located at top (PC-1) and right (PC-2). B. Representative structures obtained from cluster analysis. Most populated cluster representative is shown with red while the second and third most populated clusters representatives are depicted with yellow and green colors, respectively. The phosphorylation loops, Loop A and Loop B, located on α subunit is shown left and their counterparts on α' subunit are depicted on right.

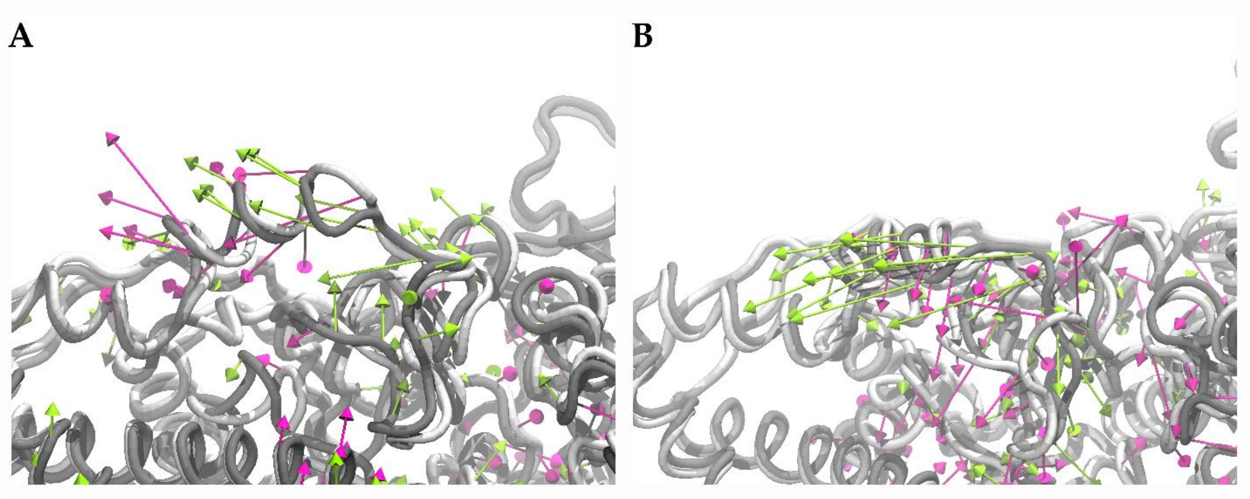

Figure S19 First principal components (PC-1) of Loop A on A. α subunit, and B. α’ subunit showing alterations of W185R (α & α’) variant (magenta) compared to WT E1 (green).

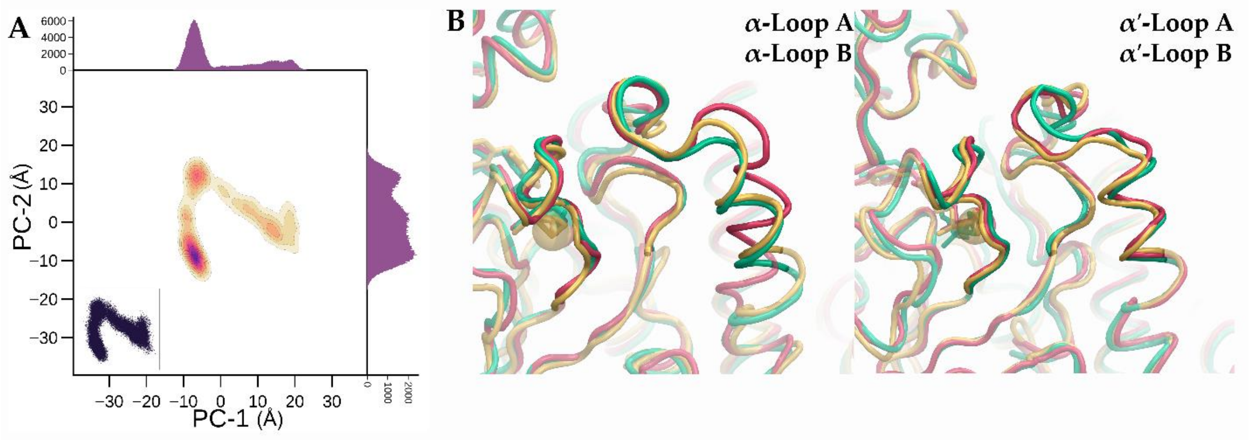

Figure S20 Principal component analysis (PCA) and cluster analysis for W185R (α & α’) variant. A. Two-dimensional density plots for the trajectory projected onto the subspace spanned by the first two principal components, PC-1 and PC-2. Scatter plot for same components is shown lower left corner while their populations are denoted with 2D histograms located at top (PC-1) and right (PC-2). B. Representative structures obtained from cluster analysis. Most populated cluster representative is shown with red while the second and third most populated clusters representatives are depicted with yellow and green colors, respectively. The phosphorylation loops, Loop A and Loop B, located on α subunit is shown left and their counterparts on α’ subunit are depicted on right.

#### 4 DYNAMIC CROSS CORRELATION ANALYSIS

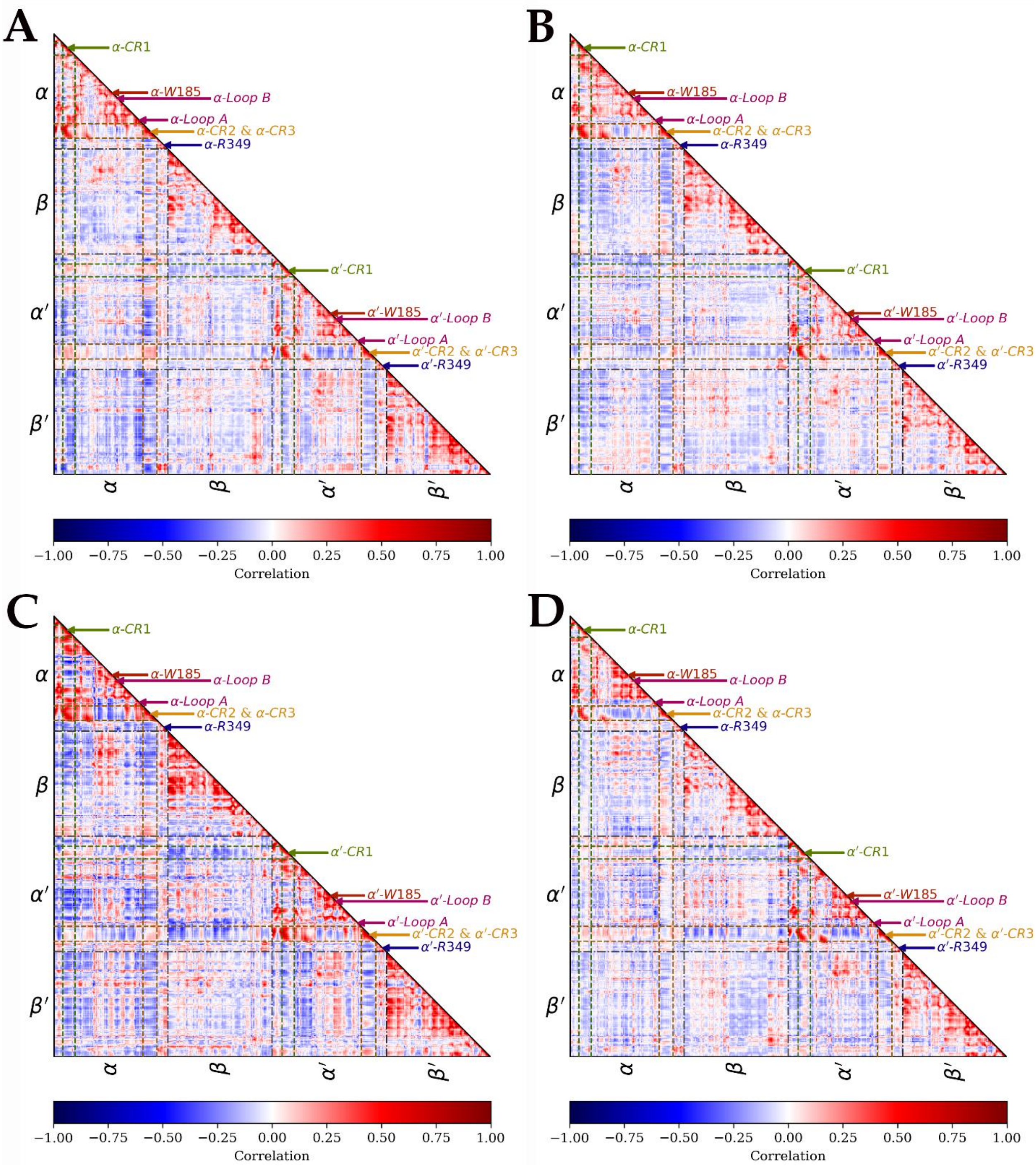

Figure S21 Lower triangular dynamic cross correlation matrices of A. R349C (α), B. R349H (α), C. W185C (α), and D.W185R (α) variants.

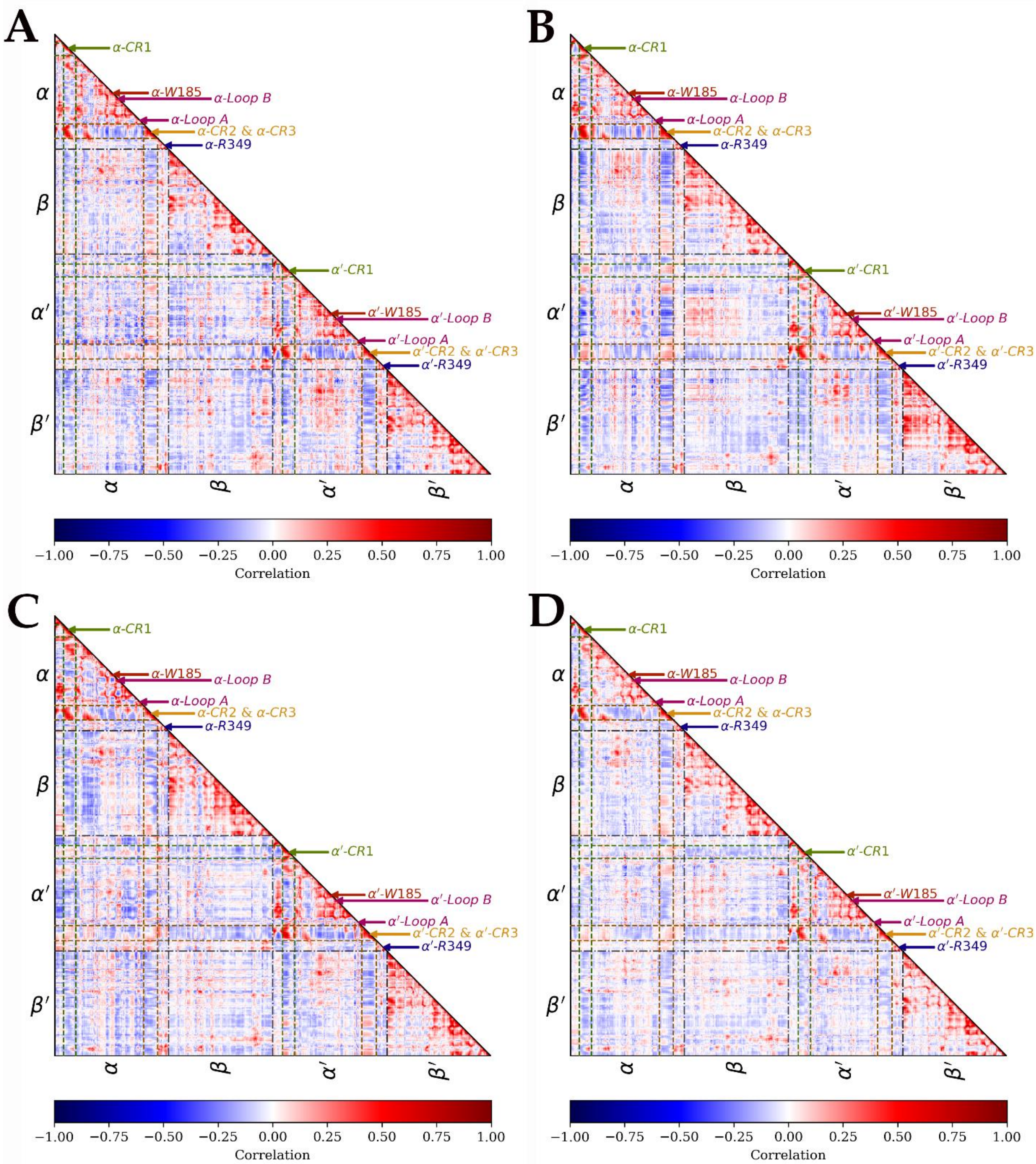

Figure S22 Lower triangular dynamic cross correlation matrices of A. R349C (α & α’), B. R349H (α & α’), C. W185C (α & α’), and D.W185R (α & α’) variants.

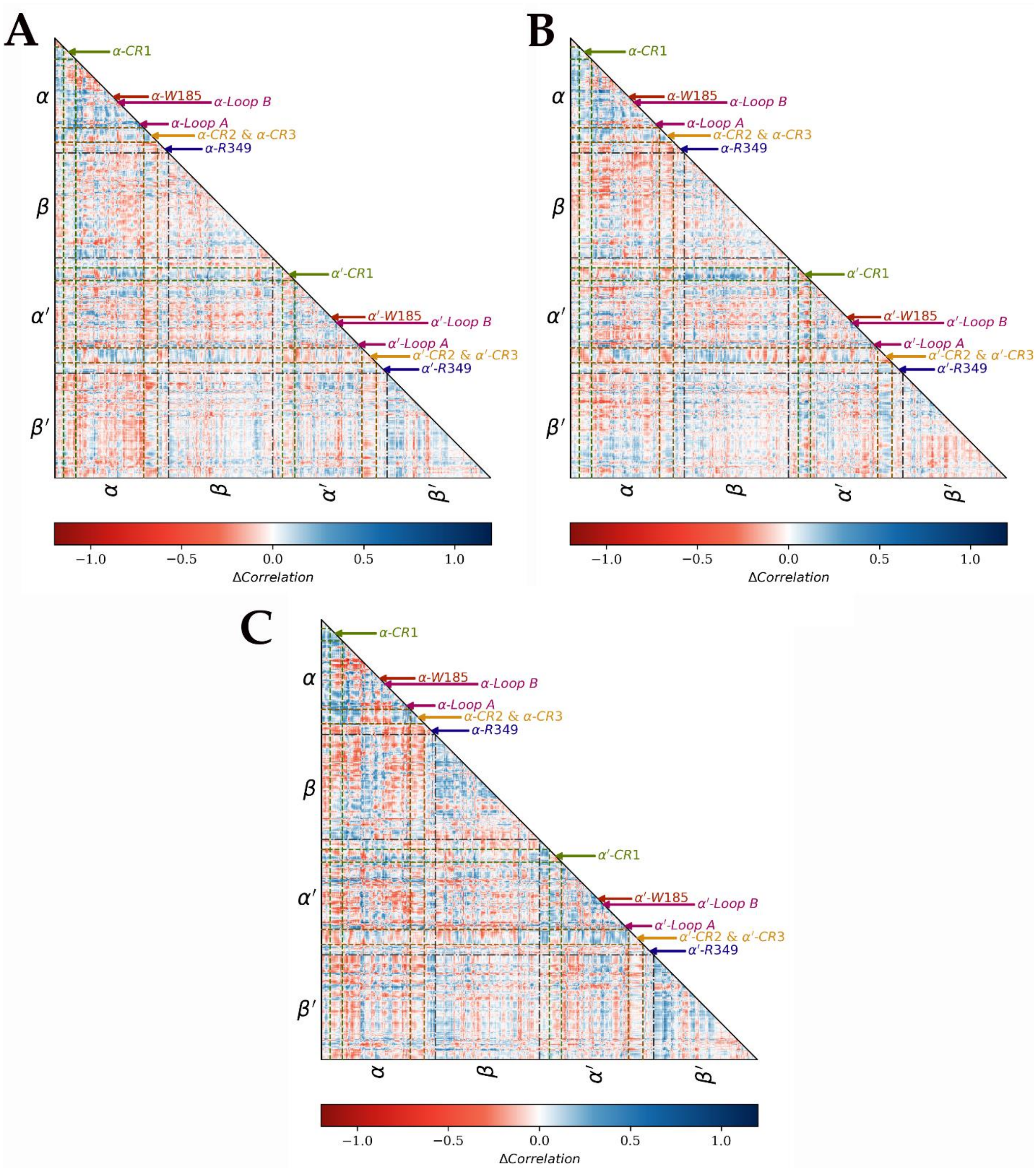

Figure S23 Lower triangular relative dynamic cross correlation matrices of A. R349C (α), B. R349H (α), and C. W185C (α) variants.

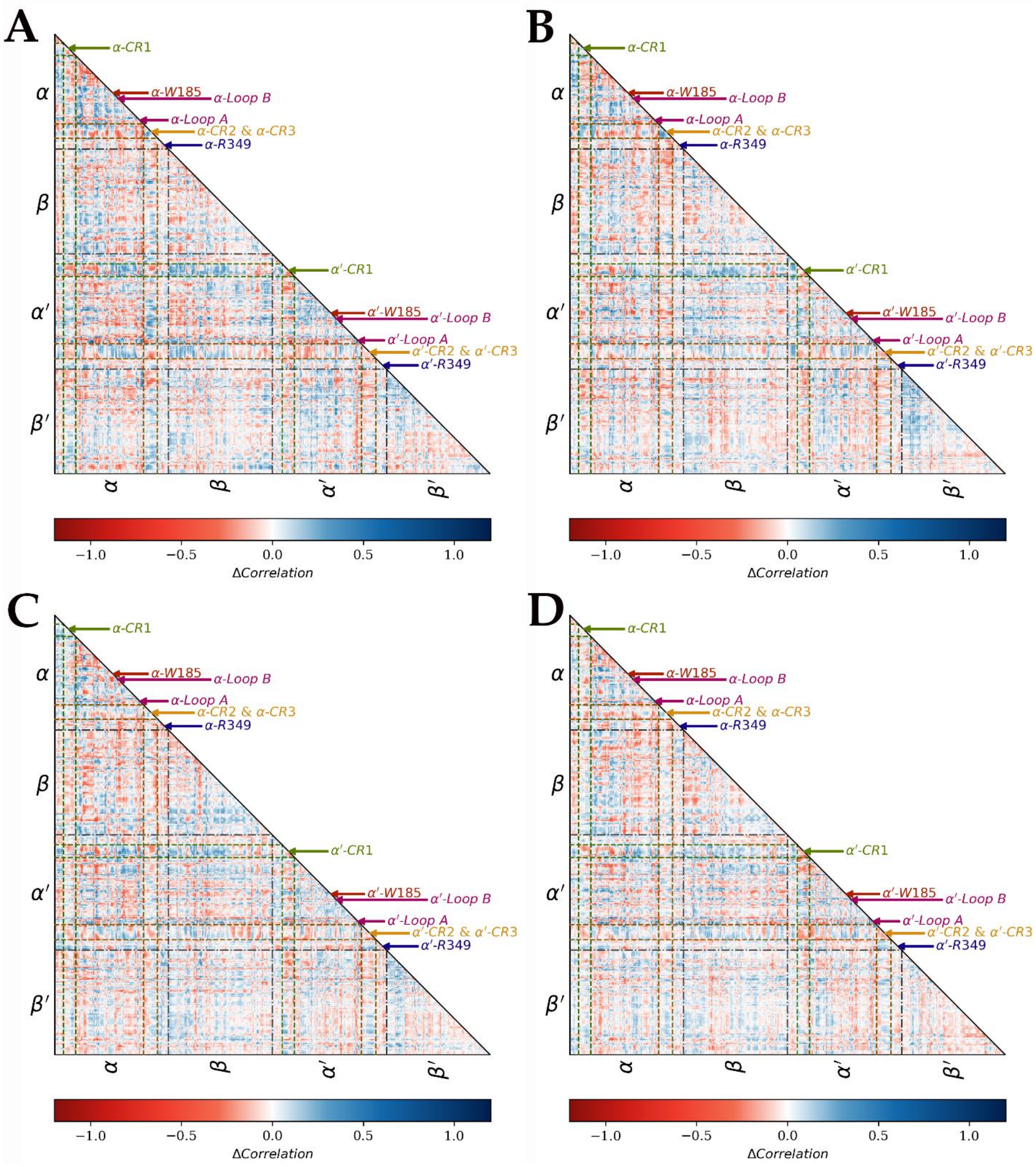

Figure S24 Lower triangular relative dynamic cross correlation matrices of A. R349C (α & α’), B. R349H (α & α’), C. W185C (α & α’), and D.W185R (α & α’) variants.

### 5 CENTRALITY ANALYSIS

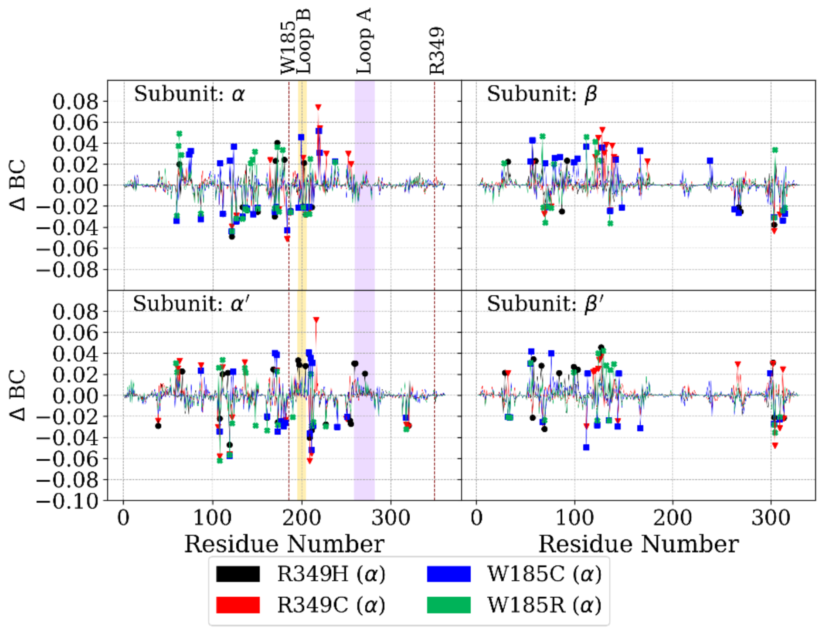

Figure S25 Relative betweenness centrality ($\Delta BC = BC_V - BC_{WT}$) of variants with mutation on α subunit. $|\Delta BC| \geq 0.02$ are shown as points.

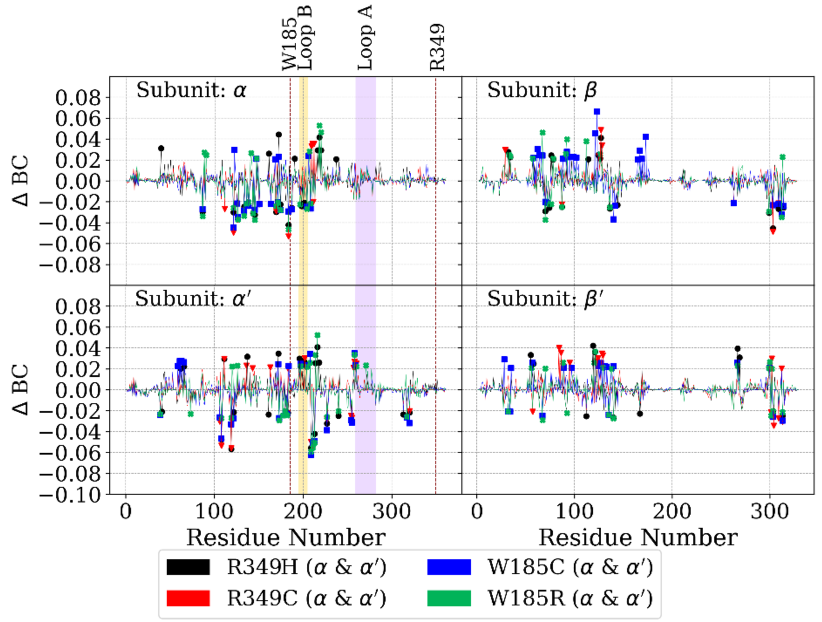

Figure S26 Relative betweenness centrality ($\Delta BC = BC_V - BC_{WT}$) of variants with mutations on α and α' subunits. $|\Delta BC| \geq 0.02$ are shown as points.

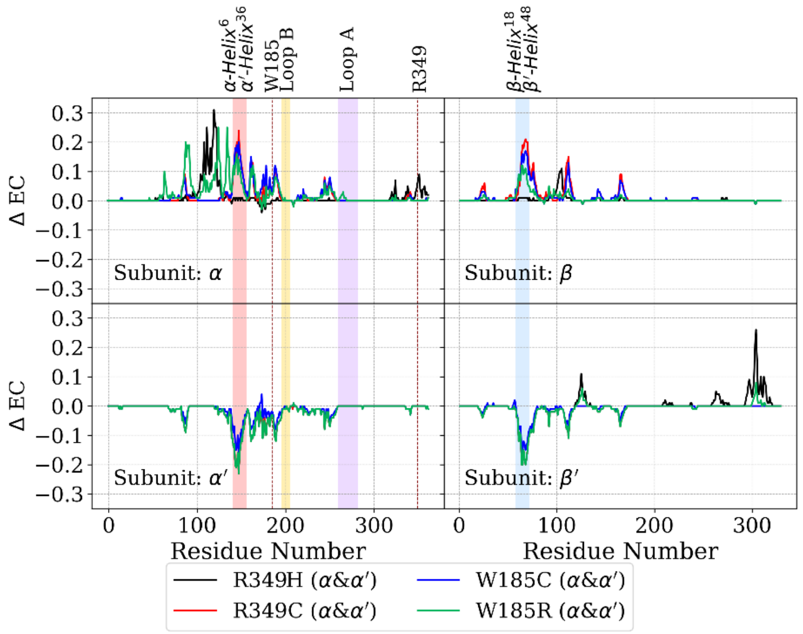

Figure S27 Relative eigenvector centrality ($\Delta EC = EC_V - EC_{WT}$) of variants with mutation on α subunit.

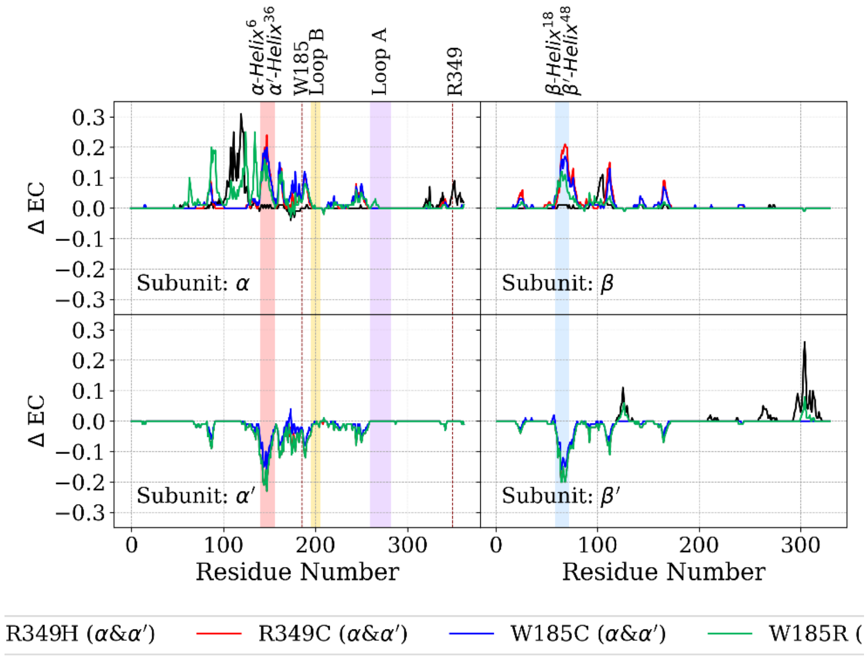

Figure S28 Relative eigenvector centrality ($\Delta EC = EC_V - EC_{WT}$) of variants with mutations on α and α’ subunits.
